# Two ancient nuclear lineages and divergent reproductive strategies drive global success of the wheat stripe rust pathogen

**DOI:** 10.64898/2026.03.18.712333

**Authors:** Jierong Wang, Yiwen Xu, Zhiru Li, Gangming Zhan, Jiasui Zhan, Zhensheng Kang, Jing Zhao

## Abstract

Understanding the mechanisms for generating and maintaining plant pathogen genetic diversity is essential for predicting disease emergence and safeguarding the food supply. Here, using population-genomic analyses of 507 global isolates of the wheat stripe rust fungus *Puccinia striiformis* f. sp. *tritici*, we uncover two nuclear lineages that diverged before modern agriculture and reinforced during wheat domestication. Distinct lineage combinations underpin three major global populations: two predominantly sympatric, homozygous lineages (nuclA–nuclA and nuclB–nuclB) and a globally dominant heterokaryotic lineage (nuclA–nuclB) of hybrid origin. This hybrid population evolves largely clonally, accompanied by recurrent somatic nuclear exchange. Identical virulence traits can arise independently via somatic nuclear exchange in clonal populations or through sexual recombination in recombining populations, revealing convergent evolutionary routes to adaptation. Together, our results provide a transformative nuclear-level perspective linking deep evolutionary history and reproductive strategy to the global success of cereal rust fungi.

## Introduction

The accelerating pace of global environmental change and rising food demand have intensified the threat to agricultural security posed by rapidly evolving plant pathogens (*1*, *2*). Understanding how pathogen diversity is generated, maintained, and reorganized is therefore both a fundamental evolutionary question and an urgent practical challenge (*3*, *4*). Among plant pathogens, cereal rust fungi in Pucciniales are particularly destructive, causing recurrent epidemics in wheat, barley, maize and other staple crops worldwide (*5*, *6*). Their extraordinary capacity to overcome host resistance, adapt to new agro-ecological environments and rapidly spread across continents makes them an ideal model for studying pathogen evolution in agricultural ecosystems (*7*, *8*).

A defining biological feature of cereal rust fungi is their heterodikaryotic urediniospore stage, in which each spore contains two genetically distinct haploid nuclei (*9*). This dikaryotic state dominates the epidemic phase and is central to their evolutionary success (*10*). The presence of two independently evolving nuclei creates unique opportunities for rapid adaptation, but the origin, differentiation and long-term dynamics of these nuclei remain poorly understood. Advances in long-read sequencing and haplotype phasing have begun to reveal extensive nuclear-level diversity in rust fungi (*11*, *12*). Studies in *Puccinia graminis* (the causal fungus of stem rust of wheat and barley) (*13*, *14*), *P. triticina* (the causal fungus of wheat leaf rust) (*15*), and *P. coronata* (the causal fungus of oat crown rust) (*16*, *17*) have documented substantial sequence divergence, structural variation and gene content differences between the two nuclei within individual isolates. Haplotype-resolved assemblies of *P. graminis* (*14*), for example, uncovered extensive genome-wide single-nucleotide polymorphisms (SNPs) (11–18/kbp) and structural variation (accounted for 6.7–8.6% of the haploid genome sizes) between coexisting nuclei. Nevertheless, the temporal trajectory of this divergence and its distribution across global populations remain largely unresolved.

Somatic hybridization and nuclear exchange have been recently documented in several cereal rust fungi (*18*). In *P*. *graminis*, haplotype-resolved genomic analyses of Ug99 lineage demonstrated that the reassortment of divergent nuclei can generate highly virulent genotypes, leading to abrupt evolutionary changes with severe epidemiological impacts (*14*). In *P*. *triticina*, population-genomic studies have shown that distinct nuclear lineages are recurrently reshuffled across global populations, indicating that ongoing nuclear exchange helps maintain population diversity even under predominantly clonal reproduction (*15*).

Despite these advances, a fundamental conceptual gap remains. Most previous studies have focused on contemporary population structure or isolated emergence events, leaving the deep evolutionary history of dikaryotic nuclei largely unexplored, particularly whether diverged nuclei constitute stable lineages that persist across regions and evolutionary timescales, and how nuclear composition interacts with reproductive strategy to shape virulence evolution. Although rust epidemics are often dominated by clonal asexual reproduction, sexual reproduction persists in regions and natural ecosystems where alternate hosts are present (*9*, *19*-*22*) and contributes to virulence variation, yet these processes have rarely been integrated into a unified nuclear-level evolutionary framework.

*Puccinia striiformis* f. sp. *tritici* (*Pst*), the causal agent of wheat stripe rust, provides an exceptional system to address these questions due to its extensive virulence diversity and mixed reproductive strategies (*23*-*26*). As a heteroecious fungus, *Pst* alternates between wheat, which supports repeated asexual reproduction, and alternate hosts (*Berberis* and *Mahonia* spp.), which enable sexual reproduction (*21*, *27*, *28*). The extended Himalayan region is regarded as a center of origin for *Pst*, where frequent sexual recombination generates high genetic diversity (*23*), whereas populations in many other regions, including North America, Europe and Australia, are dominated by clonal lineages (*29*). Despite extensive genomic analyses (*30*-*33*), how nuclear divergence, nuclear composition and reproductive strategy jointly shape the global population structure and virulence evolution of *Pst* remains unclear.

Here, we analyzed a global collection of 507 *Pst* genomes, including 11 haplotype-phased assemblies, to reconstruct the evolutionary history of this pathogen. We identify two ancient nuclear lineages that form a foundational backbone of global *Pst* populations, and demonstrate that their long-term divergence, alternative nuclear combinations, contrasting reproductive strategies, and recurrent somatic nuclear exchange, together explain the present-day global population structure and the independent emergence of key virulence traits. By integrating population genomics, haplotype-resolved comparisons and demographic inference, we provide an evidence-based evolutionary synthesis for global *Pst* emergence, with broader implications for disease surveillance based on nuclear-genotype monitoring.

## Results

### Population genomics of Chinese *Pst* isolates reveals two major clusters with contrasting reproductive signatures

To define the major population structure of *Pst* in China and assess signatures of contrasting reproductive strategies, we compiled 290 isolates sampled mainly from northwestern and southwestern China during 2005–2022 (Table S1). Among them, 258 isolates were newly sequenced and 32 were obtained from published datasets (*33*). Clean sequencing reads for each isolate were mapped to the AZ2B haplotype of *Pst* isolate AZ2 (*34*) as the reference genome, yielding an average sequencing depth of 47 × and genome coverage exceeding 97.9% for all isolates (fig. S1a). Variant calling identified 1,015,961 high-confidence variants, including 817,541 SNPs and 198,420 small insertions and deletions (indels) (fig. S1, b and c), predominantly located in upstream, downstream, and intergenic regions (Table S2). Maximum-likelihood phylogenetic analysis revealed two distinct genetic clusters, CL1 and CL2, without clear geographic or temporal separation (Fig. 1a and Table S1). CL1 comprised four subclusters (CL1a–d), with CL1a restricted to southwestern China and the remaining subclusters primarily distributed in northwestern China. CL2 comprised two subclusters (CL2a and CL2b), both distributed across northwestern and southwestern China. Principal-component analysis supported this clustering (Fig. 1b). Pairwise fixation index (*F_ST_*) values indicated pronounced genetic differentiation among subclusters (0.09–0.47), and especially between CL1 and CL2 (0.34–0.47) (fig. S2).

Phylogenetic branch lengths differed strikingly between clusters, with CL1 having uniformly short branch lengths, whereas CL2 bears longer and more heterogeneous branches (Fig. 1a), consistent with contrasting contributions of clonal propagation and recombination (*31*). Supporting this, CL1 exhibited limited diversity at the mating-type *HD* locus, with only

five allele pairs detected across subclusters, while CL2 displayed extensive diversity with 17 allele pairs, further suggesting reproductive differences between clusters. Network analysis using SplitsTree further supported these contrasting reproductive strategies (Fig. 1c and d), with CL1 having formed a star-like topology with minimal reticulation, indicative of predominant clonality; whereas CL2 exhibited extensive reticulation consistent with frequent recombination, which was confirmed by the pairwise homoplasy index (PHI) test (*p*-value ≈ 0.0; null hypothesis: clonality).

Genome-wide diversity metrics also distinguished the two clusters. CL2 exhibited much higher nucleotide diversity (π = 7.59 × 10^−2^) and Watterson’s theta (*θw* = 7.10 × 10^−2^) than CL1 (π = 1.94 × 10^−2^, *θw* = 1.89 × 10^−2^; *p* < 10^−6^, Wilcoxon rank-sum test) (Fig. 1e and Table S3). In contrast, CL1 exhibited statistically significantly higher genome-wide heterozygosity (mean = 0.58) than CL2 (mean = 0.15; *p* < 10^−6^, Wilcoxon rank-sum test) (Fig. 1f). Runs of homozygosity (ROH) analysis revealed only 1–3 short ROHs in most CL1 isolates, compared with 70–420 ROHs in CL2 (Fig. 1g). Together, these observations demonstrate that CL1 has evolved largely through clonal propagation, whereas CL2 has undergone frequent sexual recombination.

**Fig. 1.**
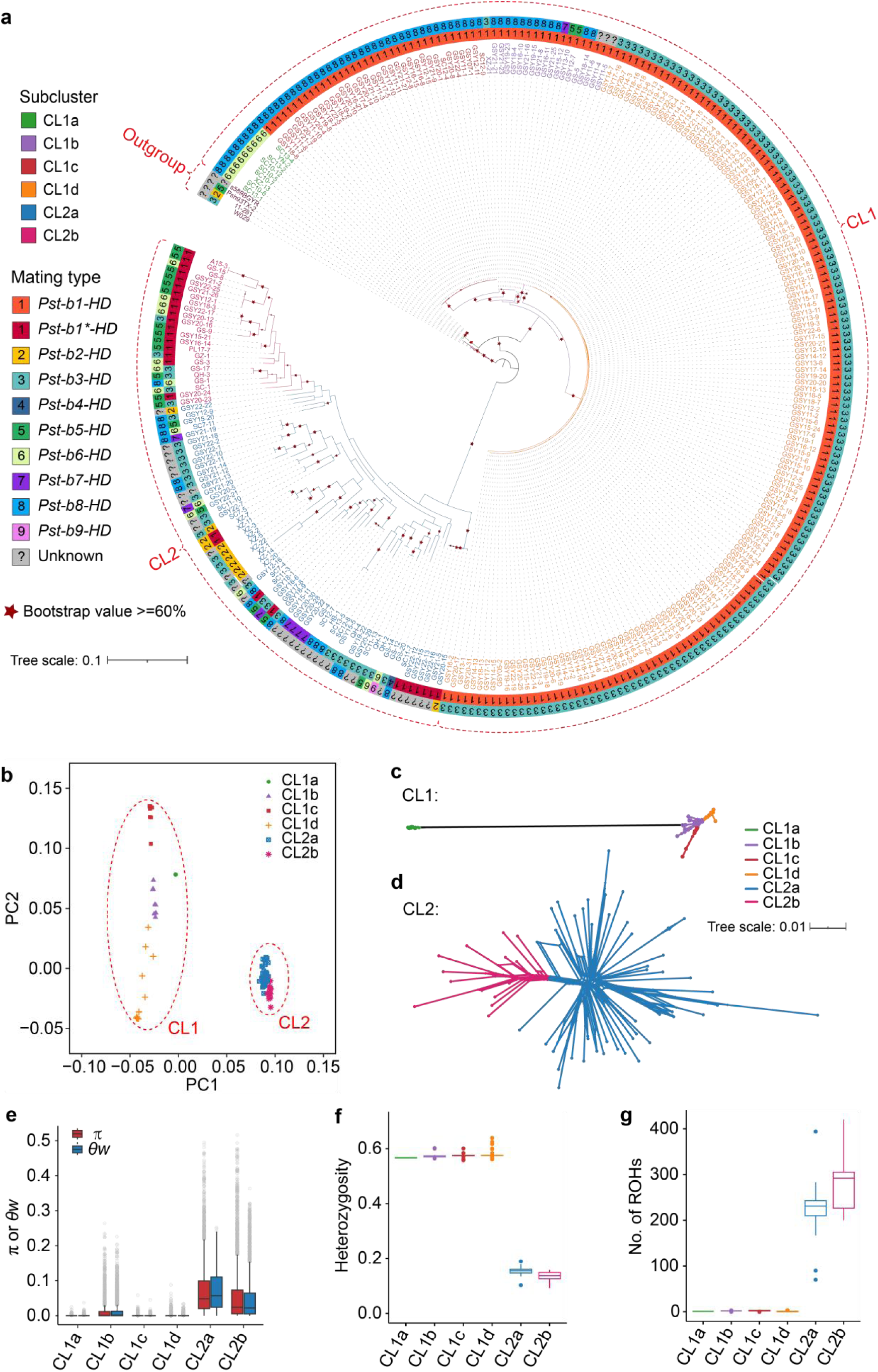
Genomic architecture and diversity of two major *Pst* clusters in China reflect contrasting reproductive strategies. **a**, Maximum-likelihood phylogenetic tree of 290 *Pst* isolates rooted using three *P. striiformis* f. sp. *hordei* isolates and a *P. striiformis* f. sp. *agropyri* isolate as outgroups. Subclusters (CL1a to CL1d and CL2a to CL2b) are indicated by colored outer rings. Mating-type alleles (*HD* genes) for each isolate are shown as paired colored bars in the outermost ring; isolates with undetermined mating types are labeled in gray. Red stars denote branches with bootstrap support ≥ 60%. **b**, Principal component analysis of 290 *Pst* isolates. **c**, Neighbor-net network for CL1 isolates based on genome-wide SNPs. **d**, Neighbor-net network for CL2 isolates based on genome-wide SNPs. **e**, Nucleotide diversity (π) and Watterson’s theta (*θw*) across *Pst* subclusters. **f**, Genome-wide heterozygosity levels for each subcluster. **g**, Number of runs of homozygosity (ROHs) per isolate. Statistical comparisons in panels **e–g** were performed using the Wilcoxon rank-sum test (*p* < 10^-6^).

### Hybrid origin and widespread nuclear exchange shape *Pst* genomic diversity

To investigate the origin of the high genomic heterozygosity in CL1, we compared observed genotype frequencies with Hardy-Weinberg expectations using *Fis* as a measure. CL1 isolates exhibited strongly negative *Fis* values (-1.02 to -0.77) (Fig. 2a), indicating substantial heterozygote excess inconsistent with long-term asexuality alone (*35*, *36*) but consistent with a hybrid origin. Site frequency spectrum (SFS) analysis further supports this hypothesis, with 27.71% of sites in CL1 being heterozygous SNPs shared across all isolates (Fig. 2b), implying that a substantial fraction of heterozygosity likely pre-dated the diversification of CL1 subclusters. Genome-wide genotype heatmaps revealed extensive heterozygosity and limited mosaic structure in CL1, whereas CL2 exhibited predominantly homozygous block-like and mosaic patterns (fig. S3), consistent with more frequent recombination in CL2. Taken together, these results suggest that CL1 likely originated through hybridization between genetically distinct lineages and subsequently diversified largely clonally, whereas CL2 shows population-genomic signatures consistent with a more recombining evolutionary history.

To validate the hybrid origin of CL1, we generated chromosome-level, haplotype-phased assemblies for five representative isolates spanning distinct clusters (Table S4). Phylogenetic analysis of haplotype sequences identified two deeply diverged nuclear lineages: nuclA and nuclB (Fig. 2c and fig. S4). NuclA was found exclusively in CL1 isolates (SC13-4, GSY17-10 and GSY16-9), with nuclB was found in both CL1 and CL2 (Fig. 2c). Pairwise identity-by-descent (IBD) analysis corroborated this structure (Fig. 2d), with nuclA haplotypes clustered together with high IBD values and were clearly separated from the nuclB cluster, confirming long-term divergence between the two nuclear lineages. Within CL1 isolates, the two haplotypes consistently exhibited minimal IBD, reflecting their marked

divergence (nuclA versus nuclB). Conversely, CL2 isolates exhibited uniformly high IBD between their two haplotypes (nuclB–nuclB). Genomic dot-plot alignments further supported the presence of two distinct nuclear lineages (fig. S5). Population-wide *k*-mer containment analysis across 290 isolates revealed a striking correspondence between nuclear composition and cluster identity. All CL1 isolates exhibited a mixed nuclA–nuclB composition, with high shared *k*-mer counts matching both nuclA and nuclB reference haplotypes, whereas CL2 isolates exhibited a nuclB–nuclB composition, characterized by strong *k*-mer similarity to all nuclB haplotypes and minimal similarity to nuclA haplotypes (Fig. 2e).

Notably, haplotypes GSY17-10A and GSY16-9A were nearly identical, differing by only ∼270 SNPs and sharing 99.99% nucleotide identity, while their companion haplotypes were highly divergent (Fig. 2c, f and Table S5). This pattern strongly indicates that these isolates share the same nuclA haplotype, consistent with somatic hybridization accompanied by nuclear exchange (*14*, *15*). Moreover, all isolates in CL1b, CL1c and CL1d shared the same mating type (*Pst-b1-HD*) and exhibited near-complete *k*-mer containment for the GSY16-9A haplotype, indicating recurrent spread of this nuclA haplotype across subclusters via repeated nuclear-exchange events (Fig. 2g and fig. S6). In contrast, several other haplotypes were largely restricted to individual subclusters or isolates (Figs. 2h–k and fig. S7, a–d). Collectively, these results demonstrate that CL1 originated through hybridization between two deeply diverged nuclear lineages, followed by long-term clonal evolution and recurrent nuclear-exchange events.

**Fig. 2.**
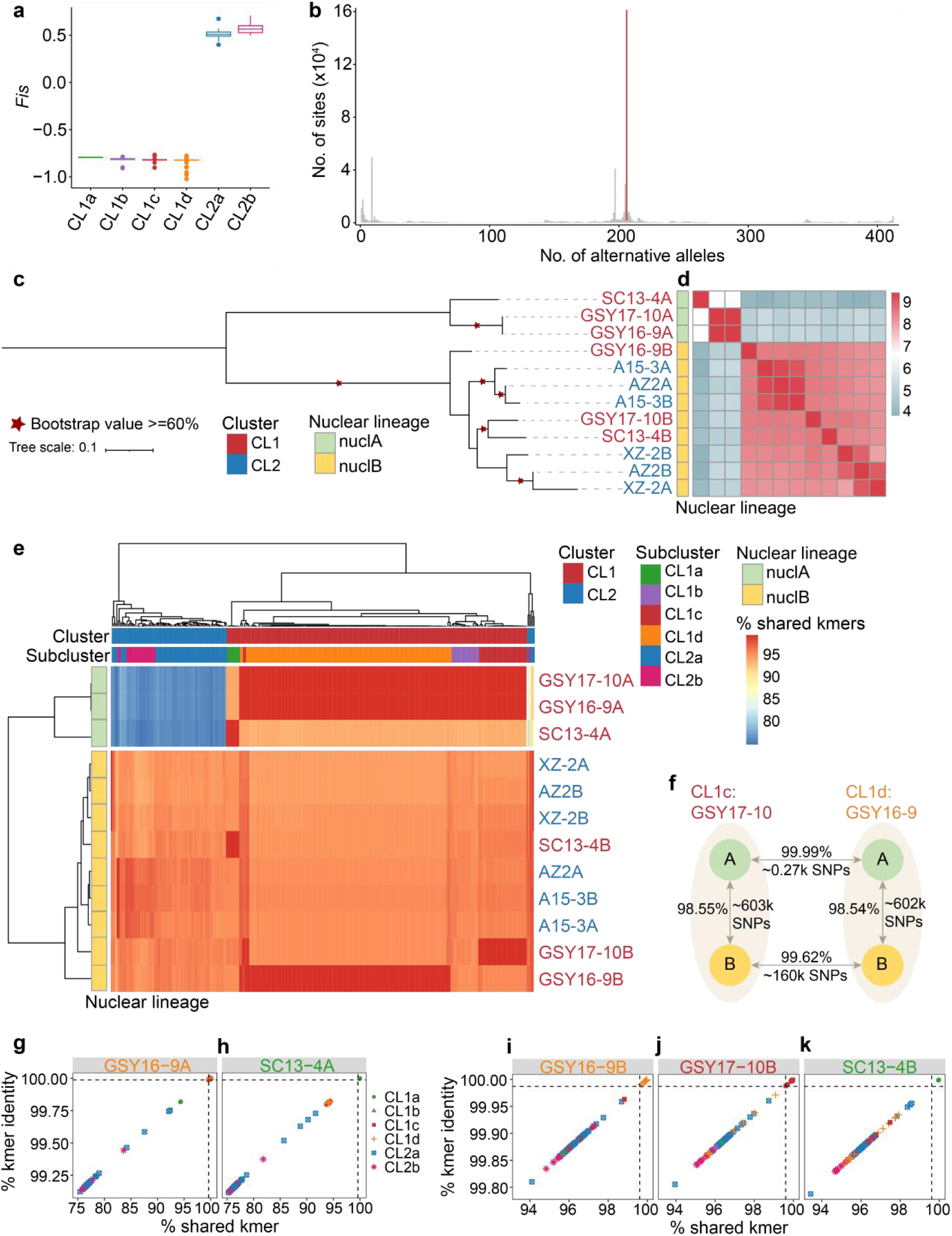
Haplotype-resolved analysis reveals two deeply diverged nuclear lineages and recurrent nuclear exchange in *Pst*. **a**, Inbreeding coefficients (*Fis*) across *Pst* subclusters. **b**, Site frequency spectrum (SFS) of 206 sampled CL1 isolates displaying the distribution of alternative allele counts (maximum = 412 alleles across 206 dikaryotic isolates). A prominent peak of shared heterozygous sites present in all CL1 isolates is highlighted in red. **c**, Midpoint-rooted maximum-likelihood phylogeny of haplotype sequences constructed from pangenome-aligned SNPs. Two deeply diverged nuclear lineages are resolved: nuclA (yellow) and nuclB (blue). Red stars mark branches with ≥ 60% bootstrap support. **d**, Pairwise identity-by-descent (IBD) matrix among haplotypes. Heatmap colors represent the log-transformed total length of IBD-sharing genomic regions between haplotype pairs. **e**, Heatmap of nuclear-lineage composition across 290 sequenced *Pst* isolates. Each row corresponds to a reference haplotype and each column to an isolate. Colors indicate the percentage of shared *k*-mers between isolate short-read data and reference haplotypes. **f**, Genomic comparison between haplotypes of isolates GSY17-10 and GSY16-9. **g–k**, Scatterplots of percent shared *k*-mers (x axis) versus percent *k*-mer identity (y axis) comparing short-read data from 290 isolates against five reference haplotypes: GSY16-9A (**g**), SC13-4A (**h**), GSY16-9B (**i**), GSY17-10B (**j**) and SC13-4B (**k**).

### Both clonal and sexual reproduction contribute to variation in *Pst* virulence

To test whether virulence patterns differ between major genetic clusters and reflect contrasting reproductive backgrounds, we compared virulence profiles across clusters. Virulence phenotypes assessed on 19 *Yr* single-gene wheat lines broadly mirrored the genetic clustering of isolates (Fig. 3a), although disparity between virulence profiles and genetic background indicated that virulence phenotypes alone are insufficient to infer population relationships. Overall virulence diversity, estimated by Nei’s diversity index (*Hs*) and the Kosman index (*K*), was higher in CL2 (*Hs* = 0.295, *K* = 0.421) than in CL1 (*Hs* = 0.220, *K* = 0.327) (Table S3). In contrast, CL1 isolates exhibited statistically significantly higher virulence complexity, defined as the number of *Yr* genes susceptible to infection (mean = 11.24), compared to CL2 isolates (mean = 9.85; *p* < 10⁻³, Wilcoxon rank-sum test) (Fig. 3b). Within clusters, CL1d and CL2b displayed the highest virulence complexity (Fig. 3c).

Analysis of virulence frequencies (VirFreq) against individual *Yr* genes revealed pronounced differences between clusters, particularly for *Yr8*, *Yr9* and *Yr76* (Fig. 3d). Compared to CL1, CL2 exhibited lower VirFreq for *Yr8* but higher frequencies for *Yr9* and *Yr76*. VirFreqs for *Yr10*, *Yr24*, *Yr26* and *Yr32* exceeded 70% in subclusters CL1d and CL2b, but remained below 20% in other subclusters. Isolates virulent to *Yr26* (V26) were first identified in southwestern China and subsequently became one of the most predominant pathotypes due to the widespread cultivation of *Yr26*-carrying wheat varieties (*37*, *38*). Given the strong genetic divergence between CL1 and CL2, the broad distribution of V26 likely reflects independent origins in the two clusters. In CL1, this pattern is consistent with virulence diversification in a predominantly clonal, heterokaryotic background, whereas in CL2 it is consistent with diversification in a more recombining population background. These results suggest two inferred evolutionary routes to similar virulence outcomes in genetically distinct *Pst* populations.

**Fig. 3.**
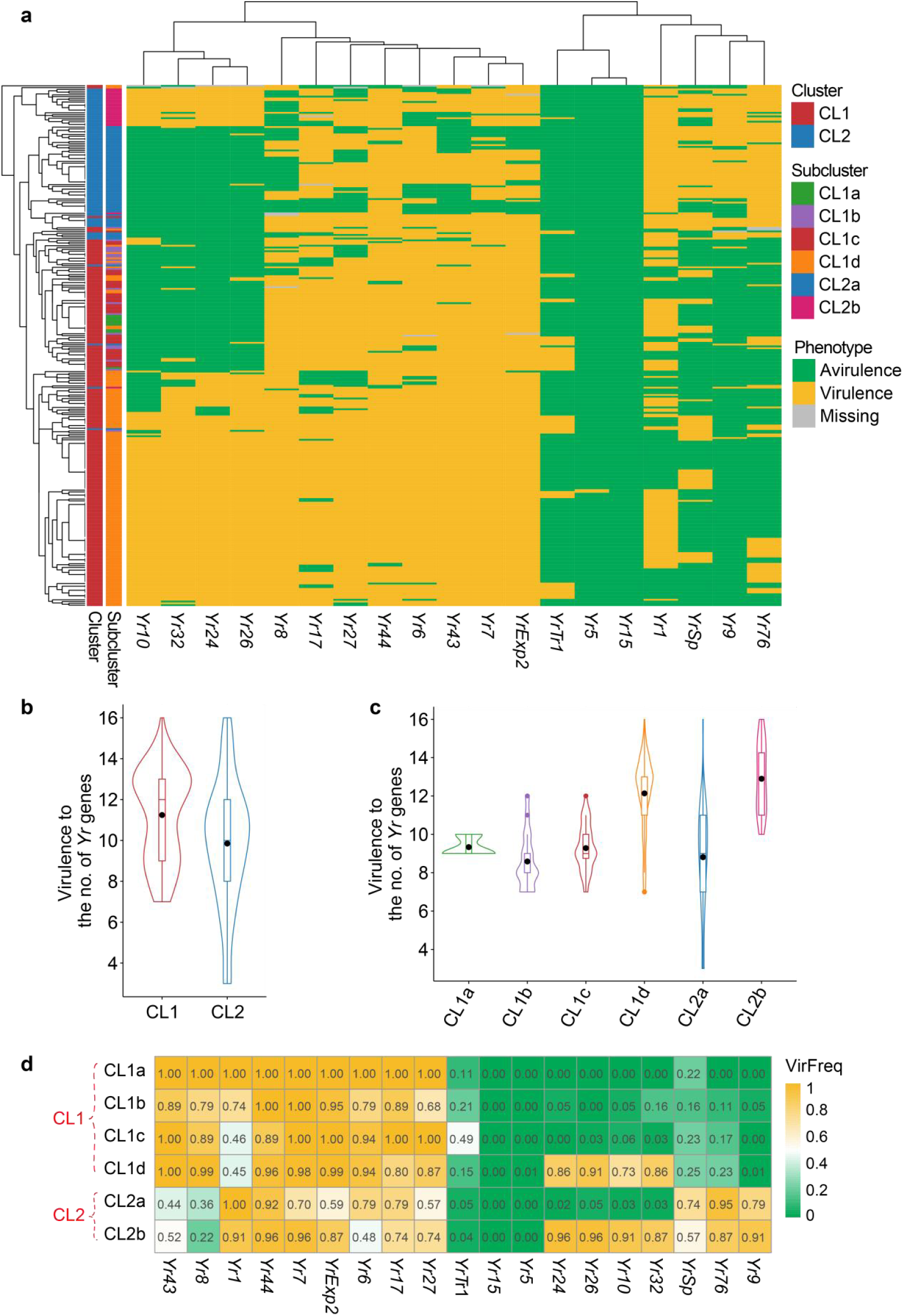
Differential virulence complexity and frequency underscore the independent evolution of *Pst* lineages. **a**, Heatmap of virulence/avirulence responses of *Pst* isolates on *Yr* single-gene wheat lines. Green indicates avirulence, yellow indicates virulence, gray denotes missing data. **b**, Distribution of virulence complexity, defined as the number of *Yr* genes to which an isolate is virulent, for isolates in CL1 and CL2. **c**, Virulence complexity across subclusters. **d**, Virulence frequency (VirFreq) of each subcluster against individual *Yr* genes. Values represent the proportion of isolates within each subcluster exhibiting virulence toward a specific *Yr* gene. Statistical comparisons in **b** and **c** were performed using the Wilcoxon rank-sum test (*p* < 10^-3^).

### The global *Pst* population structure is shaped by nuclear-lineage composition and reproductive divergence

To resolve the global population structure of *Pst* and its relationship nuclear composition and reproductive strategy, we integrated genome sequences from 217 international isolates with 290 sequences mentioned above (n = 507) (Table S6). Population-genetic analyses identified three major clusters: the previously described CL1 and CL2, and a third cluster (named CL3) predominantly distributed in southern Asia and eastern Africa (Fig. 4a and fig. S8–S9). CL2 remained largely confined to China, whereas CL1 showed broad global distribution. CL3 exhibited intermediate heterozygosity and predominantly positive *Fis* values, together with high mating-type diversity, consistent with frequent sexual reproduction (Fig. 4b and c, and fig. S8).

To resolve how nuclear-genotype composition maps onto this global population structure, we performed shared *k*-mer analyses. These revealed clear nuclear-genotype stratification across clusters, with most CL3 isolates carrying nuclA–nuclA, CL2 isolates carrying nuclB–nuclB and CL1 isolates carrying the mixed nuclA–nuclB configuration (Fig. 4d). Several isolates from Africa, Australia and the Americas shared an identical nuclA haplotype (GSY16-9A) with Chinese CL1 isolates, differing mainly in their nuclB haplotypes (fig. S9–S10). Notably, the CL1 isolate Pst239E carries the same nuclA haplotype as CL1 isolate Pst104E but a nuclB haplotype nearly identical to that of the CL2 isolate PstS7 (fig. S9), illustrating nuclear exchange between divergent lineages (*39*). These observations demonstrate that somatic hybridization and nuclear exchange have played a major role in shaping global *Pst* genomic diversity.

**Fig. 4.**
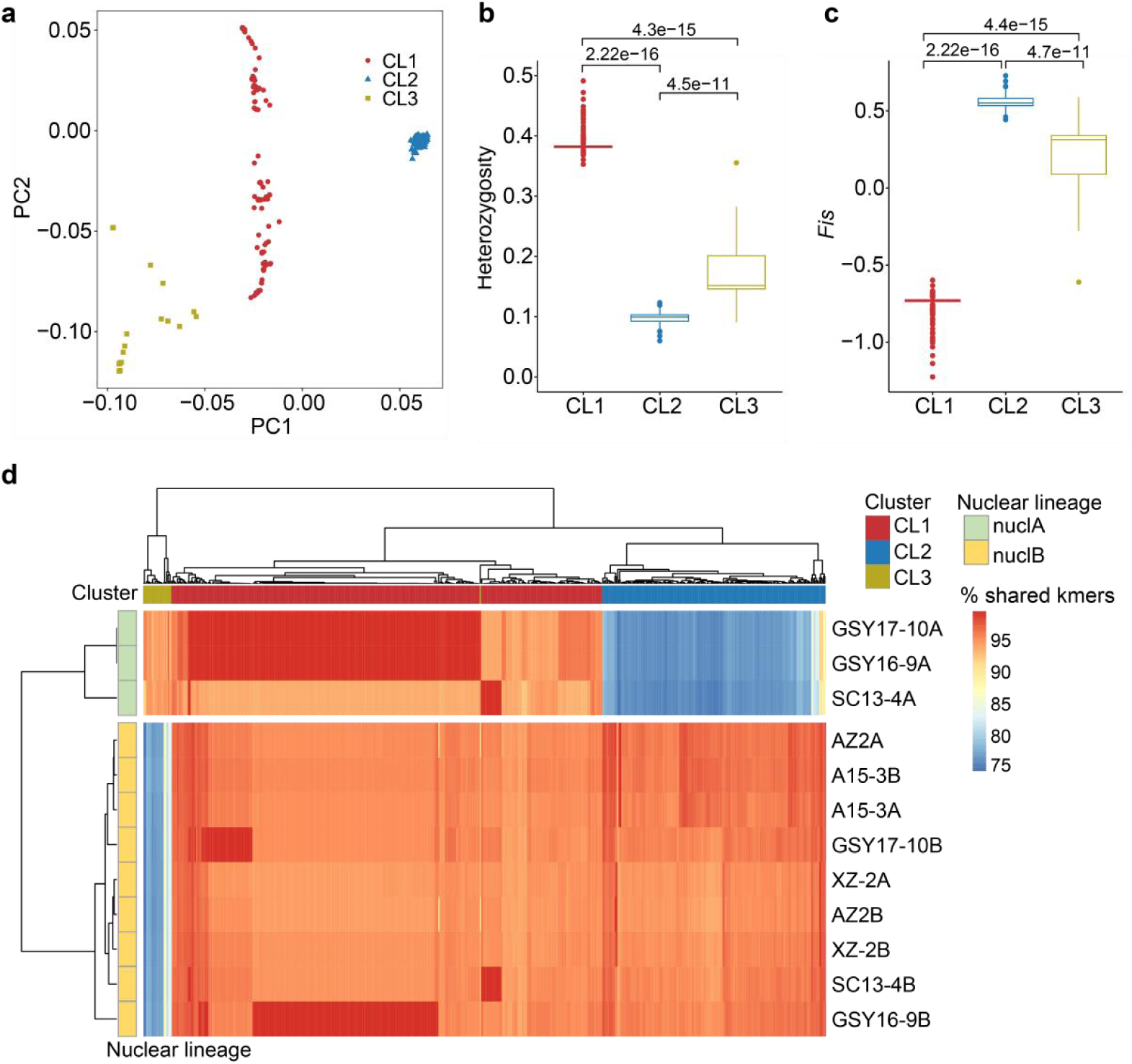
Global *Pst* isolates is defined by three discrete clusters with divergent nuclear-lineage compositions. **a**, Principal-component analysis of 507 global *Pst* isolates showing separation among the three major clusters (CL1–3). **b**, Genome-wide heterozygosity across global *Pst* clusters. **c**, Inbreeding coefficients (*Fis*) across global *Pst* clusters. **d**, Heatmap of nuclear-lineage composition among 507 global isolates. Rows correspond to reference haplotypes and columns to isolates. Colors represent the percentage of shared *k*-mers between each isolate short-read dataset and each reference haplotype.

### Phylogenetic and demographic analyses resolve ancient divergence between the two nuclear lineages of *Pst*

To refine the evolutionary relationships of the nuclear lineages that underlie global *Pst* structure, we generated haplotype-phased chromosome-level assemblies for two CL3 isolates (Table S4). The inclusion of these homokaryotic (nuclA-nuclA) backgrounds allows nuclA and nuclB to be analyzed as independent evolutionary lineages. Haplotype phylogenies incorporating CL3 assemblies confirmed nuclA and nuclB as the two principal nuclear lineages underlying global population structure (Fig. 5a). Pairwise IBD and network analyses supported deep lineage separation, with limited exceptions suggesting possible introgression

(Fig. 5b and c). In addition to the deep divergence between nuclA and nuclB, pervasive reticulation and strong rejection of clonality indicate that contemporary diversity within each lineage has been predominantly shaped by ongoing intra-lineage recombination (fig. S11a and b).

To reconstruct the divergence history of the two nuclear lineages, we performed MSMC2 analyses on representative nuclA and nuclB haplotypes. The two lineages showed broadly similar demographic trajectories, yet with consistently offset effective population sizes (*Ne*) throughout most of their histories (Fig. 5d), indicating long-term semi-independent evolution. This offset became particularly pronounced after ∼8,000–10,000 years ago, coinciding with the domestication and early global expansion of bread wheat (*40*, *41*), suggesting that ecological and host specialization and agricultural-niche formation reinforced lineage differentiation. Both nuclear lineages maintained large ancestral population sizes until

∼10^3^–10^4^ years ago, followed by a prolonged period of moderate *Ne* and a sharp recent contraction, consistent with recent clonal expansion and strong selection under agricultural intensification.

To infer how divergence and genetic connectivity between nuclA and nuclB changed through time, we performed relative cross-coalescence rate (rCCR), which revealed a parallel three-phase history (Fig. 5e). In deep time, rCCR values close to 1 indicate that the two nuclei that compose modern heterokaryotic *P. striiformis* populations originated from a single ancestral rust population during the late Pleistocene. rCCR then declined steadily through the terminal Pleistocene and early Holocene, reflecting progressive nuclear differentiation as populations became fragmented. The timing at which the two nuclei substantially diverged (rCCR ≈ 0.5) coincides closely with the origin of wheat approximately 8,000–10,000 years ago. A transient rebound in rCCR between ∼3,000 and 1,000 years ago likely reflects renewed genetic connectivity mediated by sexual recombination during which alternate hosts persisted and by somatic hybridization facilitated by expanding grasslands and early cereal cultivation. In the most recent period (<1,000 years ago), rCCR dropped sharply below 0.5, indicating strong nuclear isolation associated with predominantly asexual reproduction on cultivated wheat. Together, these results establish nuclA and nuclB as deeply diverged nuclear lineages the combinations of which, i.e. nuclA–nuclB (CL1), nuclB–nuclB (CL2) and nuclA–nuclA (CL3), underpin the global *Pst* population structure.

**Fig. 5.**
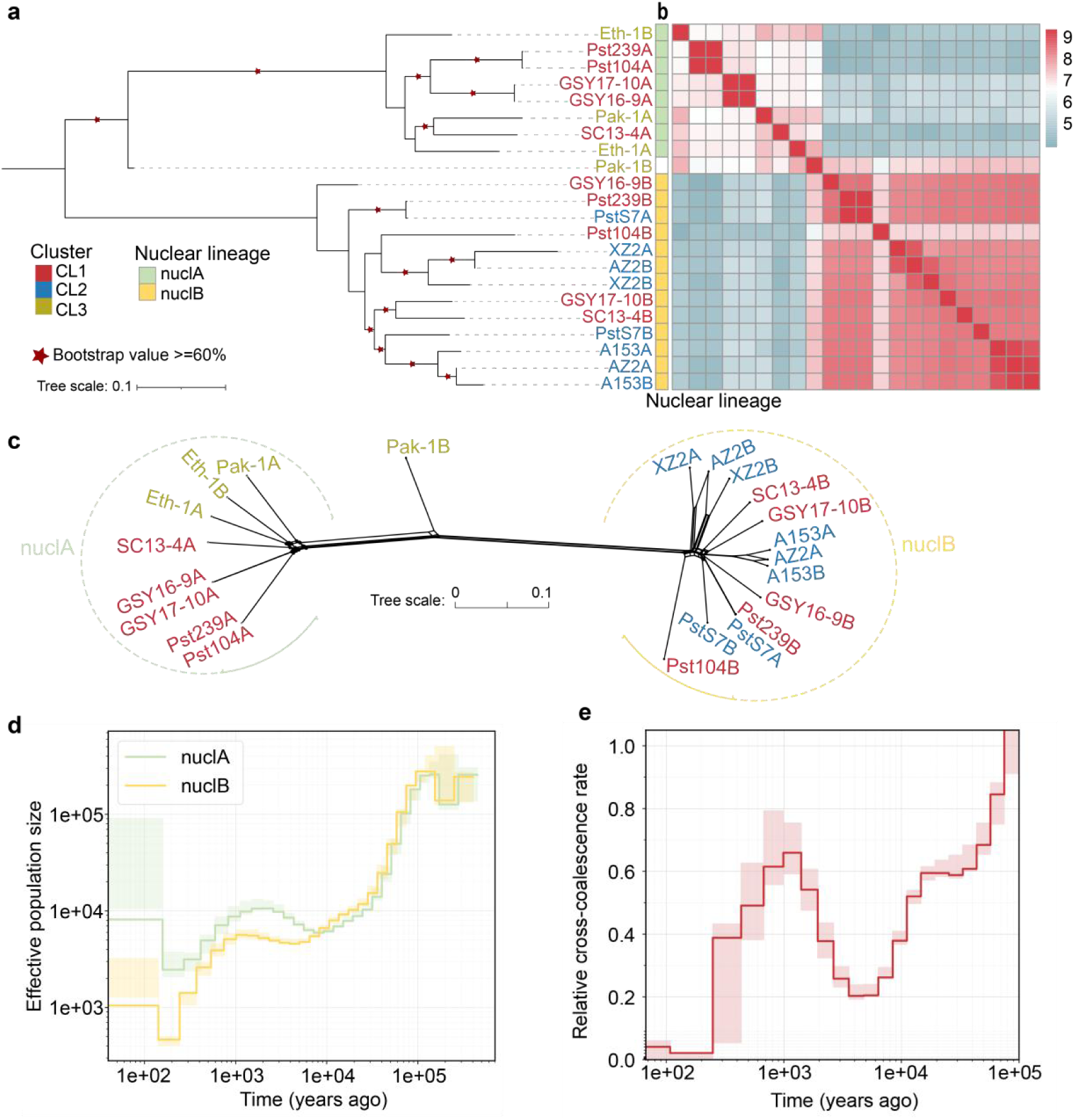
Phylogenomic structure and demographic inference uncover deep divergence between the two nuclear lineages of *Pst*. **a**, Midpoint-rooted maximum-likelihood phylogeny of haplotype sequences from representative *Pst* isolates constructed from pangenome-aligned SNPs. Red stars denote nodes with ≥ 60% bootstrap support. **b**, Pairwise identity-by-descent (IBD) heatmap of all haplotype pairs. Colors reflect the log-transformed total length of shared IBD segments. **c**, Neighbor-net network of all haplotypes. **d**, Demographic histories of nuclA and nuclB inferred using MSMC2. Lines represent estimates of effective population size (*Ne*) over time, and shaded regions indicate block-bootstrap replicates. **e**, Relative cross-coalescence rate (rCCR) between nuclA and nuclB over time. Shaded curves represent bootstrapped estimates.

### A proposed global evolutionary model of *Pst*

Based on our population genomic and demographic analyses, we propose an inference-based

evolutionary model for the global emergence and diversification of *Pst* (Fig. 6). Our data suggest that modern *Pst* populations are structured by two ancient, deeply diverged nuclear lineages that likely originated from a single ancestral population. The rCCR analyses indicate that nuclear differentiation was progressively reinforced ∼10,000 years ago. This timing is consistent with the origin and spread of bread wheat, suggesting that the creation of novel agricultural niches may have promoted host specialization. The data parsimoniously suggest that around 3,000 years ago, somatic hybridization or sexual reproduction combined these divergent nuclei, generating heterokaryotic genotypes. This dikaryotic state may have provided the genomic flexibility required for adaptation to early domesticated wheat. By approximately 1,000 years ago, the widespread cultivation of wheat likely favored predominantly asexual reproduction, leading to the stabilization and further differentiation of the two nuclei. Subsequent regional divergence, particularly in areas such as northwestern China and south Asia/east Africa, gave rise to predominantly homokaryotic nuclear configurations, nuclB–nuclB (CL2) and nuclA–nuclA (CL3). More recently, recurrent somatic hybridization between CL2 and CL3 appears to have reassembled nuclA–nuclB heterokaryons (CL1). Notably, virulence to the resistance gene *Yr26* emerged independently in CL1 and CL2, potentially through somatic hybridization-mediated variation in CL1 and sexual recombination in CL2, highlighting multiple evolutionary routes to convergent adaptive outcomes. This framework illustrates how nuclear divergence, reproductive strategy and nuclear exchange have jointly shaped the global success of *Pst*.

**Fig. 6.**
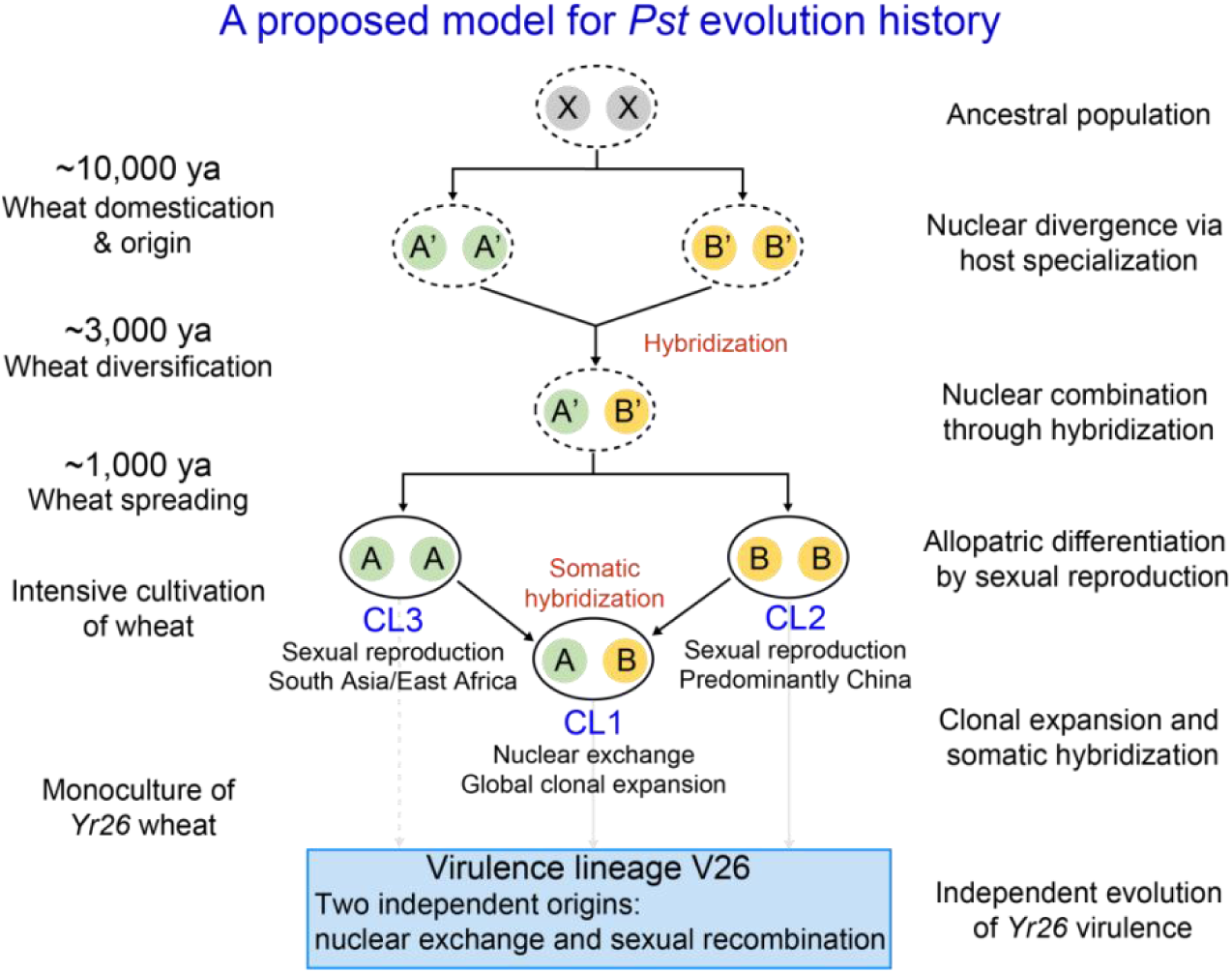
An inference-based evolutionary model of *Pst*. Two ancient nuclear lineages (nuclA and nuclB) are inferred to have originated from a single ancestral population and to have diverged in deep time, with differentiation likely reinforced during wheat domestication and early expansion (∼10,000 years ago). During a prolonged intermediate phase (schematically ∼10,000 to ∼1,000 years ago), limited genetic connectivity, including occasional hybridization/introgression, may have occurred. Within the last ∼1,000 years (schematic timescale), distinct nuclear-genotype states became predominant under contrasting reproductive regimes: sexual recombination is consistent with the predominance of nuclA–nuclA (CL3) and nuclB–nuclB (CL2), whereas hybridization between nuclA and nuclB is inferred to have generated CL1 (nuclA–nuclB). CL1 then appears to have diversified mainly through clonal propagation, with recurrent somatic hybridization and nuclear exchange contributing to high heterozygosity and broad global distribution. The model also provides a conceptual explanation for the independent emergence of virulence phenotypes (for example, V26) in genetically divergent populations. Time scales and transitions are schematic and summarize an inference-based evolutionary synthesis rather than a fully resolved historical reconstruction.

## Discussion

The rapid emergence and global spread of wheat stripe rust epidemics have long underscored the exceptional evolutionary potential of *Pst* (*42*, *43*). Yet the processes that generate and maintain this diversity, particularly the interplay among clonal propagation, sexual recombination, hybridization and the pathogen’s dikaryotic lifestyle, remain incompletely understood. By integrating 507 genomes with haplotype-phased assemblies, we support an inference-based evolutionary framework centered on two ancient and deeply diverged nuclear lineages, nuclA and nuclB. This framework helps explain major patterns of global population

structure and adaptive dynamics in *Pst*, and places long-term nuclear divergence as a major axis of pathogen diversification.

### Ancient nuclear divergence as the backbone of global population structure

Previous genomic studies have largely interpreted population structure of cereal rust fungi primarily in terms of recent clonal expansions, regional introductions and episodic sexual recombination (*13*, *30*, *32*, *44*, *45*). Our results extend this perspective by demonstrating that two ancient nuclear lineages, nuclA and nuclB, consititue a deeper evolutionary backbone underlying the major global *Pst* populations.

Demographic analyses suggest that nuclA and nuclB diverged thousands to tens of thousands of years ago from an ancestral population. The inferred timing of substantial differentiation overlaps with the origin and early spread of domesticated wheat (*40*, *41*). This temporal concordance is consistent with the hypothesis that ecological changes associated with wheat domestication and agricultural expansion may have contributed to selective environments that reinforced nuclear-lineage divergence. A similar host–pathogen association has been reported in *P. triticina*, where divergence among lineages infecting diploid, tetraploid and hexaploid wheat relatives broadly mirrors wheat domestication history (*46*). While such study has established a link between host and pathogen evolution at the population level, our analyses advances this framework by showing that long-term pathogen divergence can be retained within individual dikaryotic genomes. Rather than being fully explained as transient products of recent hybridization, nuclA and nuclB are best interpreted as ancient evolutionary entities, the divergence of which predates modern epidemics and continues to structure contemporary populations. Together, these findings suggest that cereal rust fungi may commonly preserve deep ancestral genetic structure that is subsequently reorganized and amplified by host evolution and agricultural expansion.

### Reproductive strategy links nuclear composition to regional dynamics

Nuclear composition is strongly associated with reproductive mode across global *Pst* populations. CL2 (nuclB–nuclB), predominantly found in China, and CL3 (nuclA–nuclA), concentrated in south Asia and east Africa, both bear clear signatures consistent with frequent sexual recombination. This pattern aligns with previous evidence for strong population differentiation across the Pakistan–China border (*47*) and supports a regional role for sexual reproduction in shaping diversity. Sexual reproduction likely accelerates local adaptation and

virulence diversification through meiotic recombination and reassortment of standing genetic

variation (*33*, *48*, *49*). Consistent with this interpretation, many isolates capable of completing the sexual cycle on barberry, including the European isolate DK0911, originate from the Himalayan region and belong to CL2 (*36*, *50*, *51*).

By contrast, the globally widespread CL1 cluster carries the mixed nuclA–nuclB configuration and appears to be maintained predominantly through clonal propagation. CL1 exhibits extreme genome-wide heterozygosity with complex virulence profiles that cannot be explained by long-term asexuality alone. These features are instead parsimoniously explained by a hybrid origin followed by evolutionary dynamics strongly shaped by nuclear-level processes (*52*). Supporting this model, several CL1 isolates, including isolates from Australia, show reduced teliospore production, consistent with diminished sexual capacity and long-term clonal maintenance (*23*, *53*). Beyond biological advantages, such as the preservation of heterokaryotic variation, the global distribution of CL1 likely reflects historical contingencies, including human-mediated dispersal and founder events.

### Somatic nuclear exchange as a pervasive evolutionary force

Our haplotype-resolved analyses provide genome-scale evidence that somatic nuclear exchange is a widespread and recurrent process in *Pst*. We identified multiple cases in which isolates share nearly identical haplotypes but differ substantially in their companion nuclei, a pattern consistent with repeated reshuffling of deeply diverged nuclear lineages across populations. Similar processes have been implicated in the emergence of highly virulent *P. graminis* lineages, including Ug99 (*14*), and have also been inferred from population-genomic analyses of *P. triticina* (*15*) and *P. coronate* (*16*, *17*).

Somatic nuclear exchange enables rapid genomic reconfiguration by allowing heterokaryons to combine highly diverged nuclei without meiosis (*54*). Such reassortment may generate large-effect genetic variation over short evolutionary intervals, with potential consequences for virulence and, in some cases, host range (*18*). Our results place these observations into a broader evolutionary context by suggesting that nuclear mobility operates on a background of ancient divergence and interacts with reproductive mode to generate epidemiologically important variation. More generally, in dikaryotic fungi, including rusts and many basidiomycete mushrooms, nuclear autonomy may allow selection to act at both nuclear and organismal levels, reshaping how genetic diversity is generated, maintained and expressed (*55*). Related principles have also been documented in multinucleate fungi such as *Neurospora crassa* and *Sclerotinia sclerotiorum* (ascomycete fungi) (*56*), and *Rhizophagus*

*irregularis* (arbuscular mycorrhizal fungi) (*57*, *58*), wherein genetically distinct nuclei coexist within a shared cytoplasm and contribute unequally to fitness and adaptation. These comparisons suggest that nuclear-level selection may be a widespread but still underappreciated evolutionary force across fungi.

### Multiple evolutionary pathways to virulence emergence

The apparent independent emergence of virulence to *Yr26* in CL1 and CL2 provides an example of how distinct evolutionary routes may converge on similar adaptive outcomes. The globally dominant heterokaryotic cluster CL1, virulence evolves primarily through clonal propagation but achieves high virulence complexity through somatic hybridization and whole-nucleus exchange. These processes enable rapid, large-effect genomic change without meiosis, potentially allowing clonal lineages to acquire novel virulence traits. Importantly, natural mutation accumulation still plays a critical role by fine-tuning virulence phenotypes after their initial emergence, as exemplified by *Pst* virulence evolution in Australia (*30*).

By contrast, in more sexually recombining populations such as CL2 and CL3, virulence evolution is likely shaped largely by meiotic recombination and reassortment of standing genetic variation. Sexual recombination in *Pst* can generate new virulence phenotypes and facilitate adaptation to newly deployed resistance genes (*49*, *50*, *59*). Sexual reproduction has also been reported frequently in natural populations of cereal rusts infecting barley and oat, raising the possibility that alternate hosts play a more persistent and widespread role in pathogen evolution than previously assumed (*45*, *60*, *61*). Together, these findings suggest that sexual reproduction may be a major, yet often underappreciated, contributor to virulence evolution in natural ecosystems, not only in *Pst* but potentially across cereal rust fungi more broadly.

### Implications for surveillance and disease management

Our findings have practical implications for pathogen surveillance and disease management. Traditional lineage tracking based on clonal structure or regional phylogenies may overlook nuclear-level dynamics that substantially shape *Pst* evolution (*62*-*64*). The nuclear-genotype framework established here provides a biologically meaningful basis for monitoring global populations, including routine tracking combinations of nuclA/nuclB and deriviates, their geographic movement and signatures of nuclear exchange. This could strengthen early warning by prioritizing newly detected nuclear combinations, rapid spread of virulence

phenotypes against widely deployed resistance genes and cross-regional movement of high-risk lineages.

Surveillance design should also be adapted to regional reproductive context. In regions with frequent sexual recombination, denser temporal sampling may improve detection of newly generated virulence combinations. In predominantly clonal regions, surveillance may be more informative when focused on dispersal, persistence and reassortment of successful heterokaryotic lineages. In both settings, integrating genomic surveillance with virulence phenotyping and host cultivar deployment data, together with standardized cross-regional data sharing, should improve risk assessment and forecasting. More broadly, the inference that nuclear-lineage divergence predates modern agriculture highlights a central surveillance challenge: adaptive potential may be underestimated when monitoring focuses only on recent population processes.

### Limitations and future directions

While our study offers valuable insights into the evolutionary dynamics and global population structure of *Pst*, several limitations must be addressed. First, the 507 genome sequences analyzed may not fully represent populations in key wheat-growing regions, such as the Middle East, North Africa, Central Asia, and parts of the Mediterranean. These regions are historically important for wheat domestication, diversification, and pathogen emergence, and may harbor unsampled or transitional nuclear lineages. Second, our analyses focus exclusively on *P. striiformis* f. sp. *tritici*. Inclusion of closely related formae speciales and host-associated lineages would provide critical evolutionary context. Third, since timing from demographic inference remains subject to parameter uncertainties, calibration using historical isolates, herbarium specimens, or ancient DNA may strengthen evolutionary timescale reconstruction.

Future work integrating broader geographic sampling, cross-host comparative genomics, and functional assays of nucleus-specific virulence determinants will be crucial to dissect how nuclear composition, reproductive mode, and host ecology jointly shape long-term pathogen evolution.

## Conclusion

In conclusion, our study highlights a broader evolutionary theme in cereal rust fungi. Ancient genetic structure rooted in the dikaryotic lifestyle may be subsequently reshaped by host domestication, reproductive strategy and nuclear exchange. By placing nuclear evolution at the center of rust biology, we provide a revised evolutionary framework and a foundation for anticipating pathogen trajectories in increasingly managed and interconnected agricultural ecosystems.

## Materials and Methods

### Sample collection, purification and multiplication

Isolates of *Pst* were collected from northwestern and southwestern China from 2005 to 2022 (Table S1). The susceptible wheat (*Triticum aestivum*) cultivar Mingxian 169 was planted in 10 × 10 cm pots with 10 seeds per pot. After the first leaves fully expanded, seedlings were thinned to 4–5 plants per pot. A single urediniospore from each sample was picked under a microscope using a fine glass needle and inoculated onto one leaf, and leaves in each pot received spores from only one isolate. Inoculated seedlings were incubated in darkness at 10°C and 100% humidity for 24 h, then transferred to a growth chamber programmed for 8 h darkness (10°C) and 16 h light (15°C). Sporulation began approximately 10 d after inoculation. Pots were thinned to one sporulating plant, and plastic dividers were used to prevent cross-contamination. Urediniospores from each individual plant were harvested separately and stored at 4°C and then propagated on Mingxian 169 to obtain an adequate quantity for whole-genome sequencing and virulence assays.

### Whole-genome resequencing and variant calling

Genomic DNA was extracted from urediniospores using a cetyltrimethylammonium bromide (CTAB)-based method (*65*). DNA quality was assessed on 1% agarose gels to check for degradation and contamination. Sequencing libraries (insert size ∼350 bp) were prepared for the Illumina HiSeq platform (150-bp paired-end reads). Raw read quality was evaluated with FASTQC v0.11.4 (http://www.bioinformatics.babraham.ac.uk/projects/fastqc/), and low-quality reads were trimmed using Trimmomatic v0.36 (*66*).

Reads were mapped to the AZ2B haplotype of *Pst* isolate AZ2 (*34*) using BWA-MEM (*67*). BAM files were sorted and deduplicated using SAMtools v1.8 (*68*). Mapping statistics were assessed using QualiMap v2.2 (*69*). Variants were called using GATK HaplotypeCaller v4.4 (*70*). SNPs were filtered using GATK VariantFiltration with specific thresholds (--filter-expression FS >10.0 || MQ <40.0 || ReadPosRankSum <-8.0 || SOR >3.0 ||

MQRankSum <-12.5 || QD <2.0 --cluster-size 3 --cluster-window-size 10). Indels were filtered with the thresholds (QD <2.0 || FS >200.0 || ReadPosRankSum <-20.0). Additional filtering using VCFtools v0.1.14 (*71*) retained biallelic variants with the parameters (--max alleles 2 --min-alleles 2 --min-meanDP 5). Only indels of 1–50 bp were used in downstream analyses. Functional annotation was performed with SnpEff v4.3t (*72*).

### Population structure and phylogenomic inference

Because dense SNP datasets are computationally demanding for RAxML (*73*), we thinned SNPs with < 20% missing rate using PLINK v1.90 (--thin 1000) (*74*). Maximum-likelihood trees were reconstructed using RAxML v8.2.9 (*73*) with 100 bootstrap replications under the GTRCAT model (-f a -m GTRCAT -# 100), and the final tree was produced by integrating bootstrap support with the best-scoring topology (-f b -z -t -m GTRCAT). Principal-component analysis was performed with GCTA v1.25.3 (*75*) using biallelic SNPs filtered for a minor allele frequency (MAF) > 0.05 and no missing data.

For phylogenetic network analysis, SNPs with MAF > 0.05 and no missing data were converted from the VCF to NEXUS format using VCF2PHYLIP v2.0 (https://github.com/edgardomortiz/vcf2phylip). Neighbor-net networks and PHI tests for recombination (*76*) were performed using SplitsTree v6.3.32 (*77*).

### Mating-type analysis

Mating-type structure of *Pst* was analyzed as previously described (*31*). Allele-specific *k*-mers were indexed using Sourmash (*78*) (-p k=21,scaled=10), and allele presence in raw reads was assessed with Sourmash containment (--threshold 0). Isolates with >80% *k*-mer identity for an allele were considered positive. Additionally, reads were mapped to known mating-type alleles via BWA-MEM (*67*), and coverage was summarized using samtools depth as an independent confirmation.

### Genetic diversity, *F*is, ROH and SFS analyses

Based on biallelic SNPs with MAF > 0.05 and no missing data, VCF2PHYLIP v.2.0 (https://github.com/edgardomortiz/vcf2phylip) was used to convert SNPs in the VCF format to PHYLIP format. Nucleotide diversity (*π*) and Watterson’s theta (*θw*) were calculated using VARISCAN v.2.0 (*79*). Heterozygosity was estimated as the proportion of heterozygous SNPs. Genome-wide differentiation (*F*_ST_) in 20-kb windows was estimated using pixy v1.2.10 (*80*), excluding windows with < 10 SNPs. *F*is was calculated using VCFtools v0.1.14 (--het) (*71*). Runs of homozygosity (ROH) were detected using PLINK with specific parameters (--maf 0.05 --homozyg-density 50 --homozyg-gap 10 --homozyg-kb 40 --homozyg-snp 50 --homozyg-window-het 1 --homozyg-window-missing 5--homozyg-window-snp 50). The site frequency spectrum (SFS) was calculated using Pop-Con (https://github.com/YoannAnselmetti/Pop-Con) with default settings. Differences among populations were tested using Wilcoxon rank-sum tests in R.

### Genome assembly and haplotype comparison

High-molecular-weight DNA from *Pst* isolates SC13-4, GSY17-10, GSY16-9, A15-3, XZ-2, Eth-1 and Pak-1 was sequenced using PacBio HiFi technology (coverage ∼79–474×; Table S. 7). Hi-C libraries for GSY17-10, GSY16-9, Eth-1 and Pak-1 were prepared using *Dpn*II and sequenced on an Illumina NovaSeq system (∼7.38–9.13 Gb; Table S. 8). For these four isolates, PacBio HiFi reads and Hi-C reads were assembled using Hifiasm v0.24.0 (*81*) in the Hi-C mode. For SC13-4, GSY16-9 Hi-C reads were used to scaffold its PacBio assembly. All assemblies were scaffolded with RagTag v2.1.0 (*82*) using the AZ2 genome as a reference(*34*). Assembly completeness was assessed with BUSCO v3.0.2b (*83*) using the Basidiomycota Odb10 dataset (https://busco.ezlab.org/frames/fungi.htm), and telomeric sequences were identified following the approach described previously (*14*).

The cactus-pangenome pipeline from Cactus v2.6.7 (*84*) was applied to all haplotypes, and SNPs were called using the haplotype AZ2B (*34*) as the reference. SNPs were filtered using VCFtools v0.1.14 (*71*) with the criteria (--max alleles 2 --min-alleles 2 --max-missing 0.8). VCF files were converted to PHYLIP and NEXUS formats with VCF2PHYLIP v2.0 (https://github.com/edgardomortiz/vcf2phylip) for phylogenetic analysis. Maximum-likelihood trees were inferred using RAxML v8.2.9 (*73*) as described above. Neighbor-net networks were generated using SplitsTree v6.3.32 (*77*), and the PHI test (*76*) was used to assess evidence for recombination relative to the null hypothesis of clonality. Pairwise IBD segments among haplotypes were identified with hap-ibd v1.0 (*85*) using biallelic SNPs with no missing data.

Haplotype dot-plot alignments were generated using D-GENIES (*86*) with Minimap2 v2.28 (*87*) as the aligner. Haplotypes from isolates GSY17-10 and GSY16-9 were additionally compared using MUMmer v4.0 with nucmer (--maxmatch -c 100 -l 50), and alignment metrics were summarized using the MUMmer dnadiff module (*88*).

### Population screening for haplotype containment

The *Pst* isolates sequenced with Illumina short reads were screened against each haplotype using Mash v2.3 (*89*). These haplotypes were processed with mash sketch (-s 500,000 -k 32) and mash screen was run comparing the Illumina data against the haplotype sketch file. Shared *k*-mer counts were visualized using the pheatmap package in R (https://CRAN.R-project.org/package=pheatmap) with the default clustering method.

### Demographic-history reconstruction

Multiple Sequentially Markovian Coalescent (MSMC2) v2.1.4 (*90*) was used to infer the demographic history of nuclear lineages nuclA and nuclB. Coalescence analyses use a backwardin-time algorithm, starting from the current generation and randomly building back the lineage of genomes. Demography analysis and cross-coalescence analysis were conducted using six haplotypes for each nuclear lineage (nuclA: Eth-1A, Eth-1B, Pak-1A, GSY16-9A, Pst104A and SC13-4A; nuclB: GSY16-9B, Pst104B, SC13-4B, GSY17-10B, PstS7A and PstS7B). To quantify uncertainty, we performed block bootstrapping using multihetsep_bootstrap.py (-n 40 -s 1000000 --chunks_per_chromosome 10 --nr_chromosomes 18 --seed 2025) (https://github.com/stschiff/msmc-tools). A mutation rate of 2.0×10^−8^ per site per generation was used (*32*).

### Virulence testing

Nineteen *Yr* single-gene wheat lines, each carrying a single *Yr* gene, were used for virulence assays. Fresh urediniospores were inoculated onto expanded first leaves, and plants were incubated under the same conditions used for isolate multiplication. Infection types were recorded 18 d post-inoculation on a 0–9 scale (*91*). Types 0–6 were considered avirulent, and 7–9 virulent. Phenotypic diversity was quantified using Nei’s diversity (*Hs*) and the Kosman index (*K*) with the Virulence Analysis Tool (VAT) software (*92*). Virulence complexity was defined as the number of differential lines on which an isolate was virulent. Statistical differences among clusters were tested using Wilcoxon rank-sum tests in R.

## Acknowledgments

We thank Professor Qiuzhen Jia (Gansu Academy of Agricultural Science) and Professor Yunliang Peng (Sichuan Academy of Agricultural Science) for providing the pathogen samples. We also acknowledge the High-Performance Computing (HPC) of Northwest A&F University for providing computing resources.

## Author contributions

Z.S.K. and J.Z. designed and supervised the study and interpreted all data sets. G.M.Z. assisted with experimental design. J.R.W., Y.W.X. and Z.R.L. performed sample purification, multiplication and virulence testing. J.R.W. and J.Z. analyzed and interpreted all data sets and wrote the manuscript. J.S.Z. offered suggestions for the study and revised the manuscript. All authors contributed to manuscript revision, read and approved the submitted version.

## Data and Materials Availability

All data needed to evaluate the conclusions in the paper are present in the paper and/or the Supplementary Materials. Raw whole-genome sequences of new isolates are available in the National Center for Biotechnology Information (NCBI) database (BioProject ID PRJNA1209814). The PacBio HiFi sequencing data and Hi-C sequencing data have been submitted to NCBI (BioProject IDs PRJNA1026731, PRJNA1026730, PRJNA1385444, PRJNA1385513, PRJNA1385636, PRJNA1385708, and PRJNA1385783). Genome assemblies are available on Zenodo (https://doi.org/10.5281/zenodo.17958532).

## Supplementary Figures

**fig. S1.**
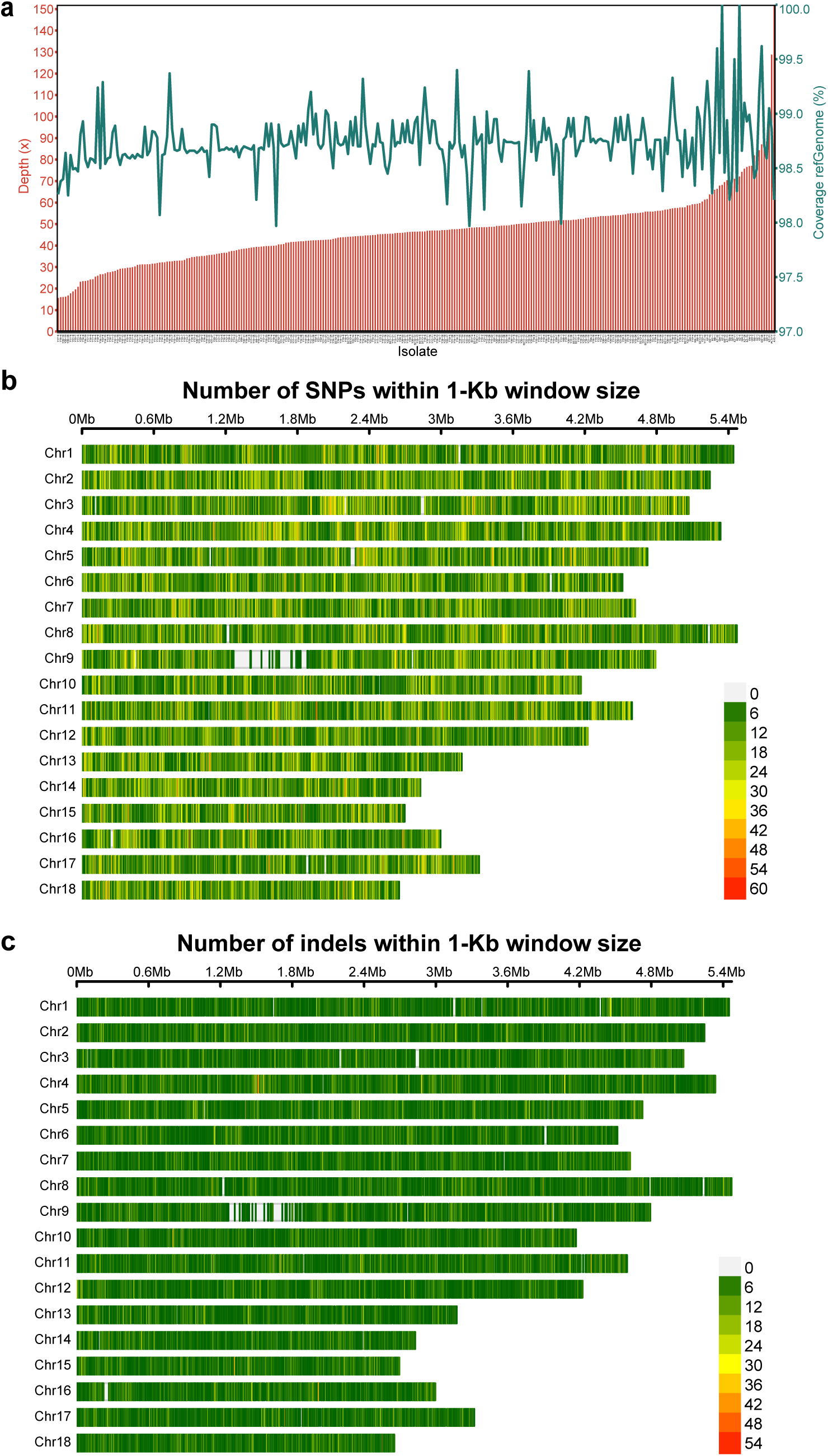
Mapping statistics and variant density across the *Pst* genome. **a**, Sequencing depth (left axis) and reference-genome coverage (right axis) for each sequenced *Pst* isolate. Bars indicate coverage percentage and the overlaid line shows mean sequencing depth. **b**, Genome-wide SNP density across the 18 chromosomes of the *Pst* reference genome, calculated in 1-Kb non-overlapping windows. Heatmap colors represent SNP counts per window. **c**, Genome-wide indel density across the *Pst* reference genome, calculated in 1-kb windows. Heatmap colors represent the number of indels per window.

**fig. S2.**
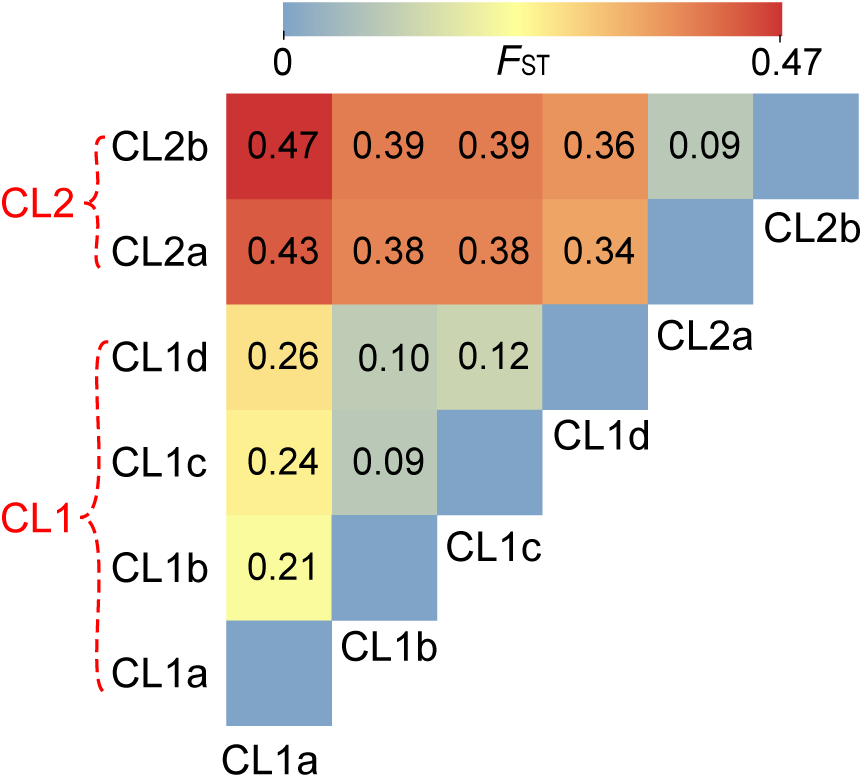
Pairwise *F*_ST_ estimates support strong genetic differentiation between *Pst* clusters. Heatmap showing pairwise *F*_ST_ values between all identified subclusters. Warmer colors indicate higher levels of genetic differentiation. Subclusters belonging to CL1 and CL2 are annotated on the left.

**fig. S3.**
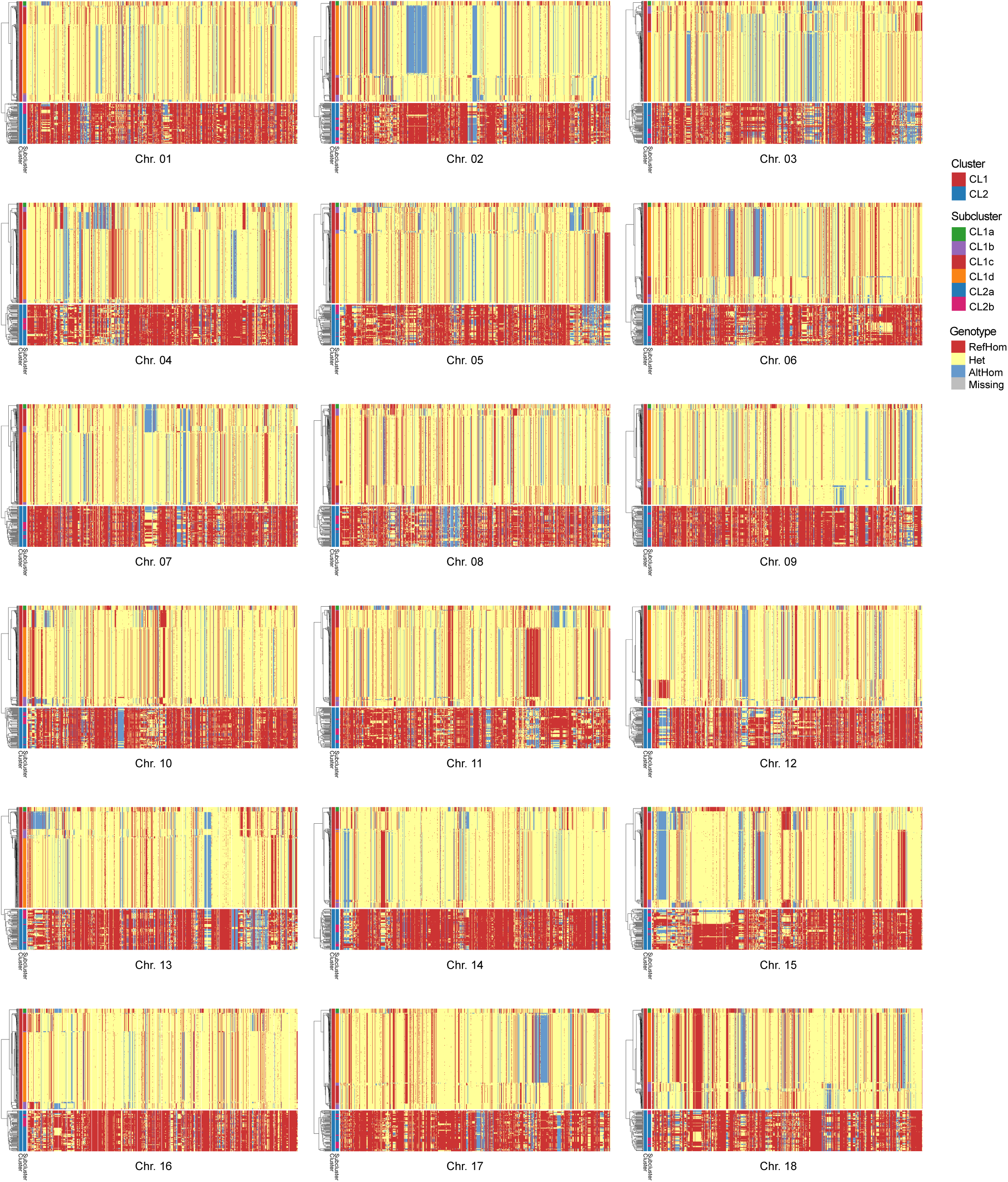
Chromosome-wide SNP genotype heatmaps show contrasting heterozygosity and mosaic patterns across Chinese *Pst* isolates. Genotype heatmaps for all 18 chromosomes. Each column represents a SNP position along the chromosome, and each row corresponds to a *Pst* isolate. SNP genotypes are color-coded as follows: red, reference homozygous; blue, alternate homozygous; yellow, heterozygous; gray, missing data.

**fig. S4.**
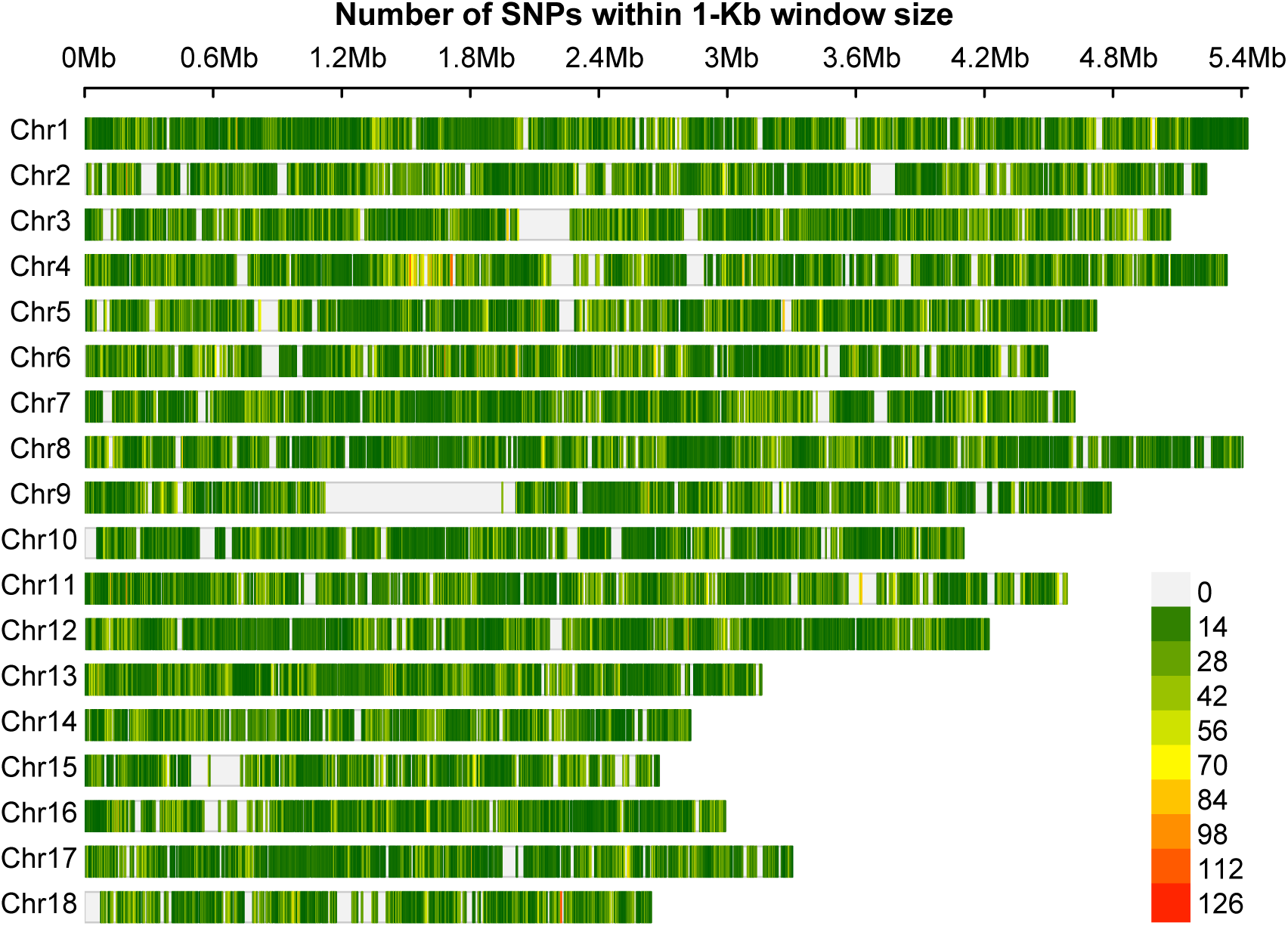
Genome-wide SNP density across *Pst* haplotypes. Genome-wide distribution of SNPs across the 18 chromosomes of the *Pst* reference genome calculated in 1-Kb non-overlapping windows for all haplotypes. Heatmap colors indicate SNP density per window.

**fig. S5.**
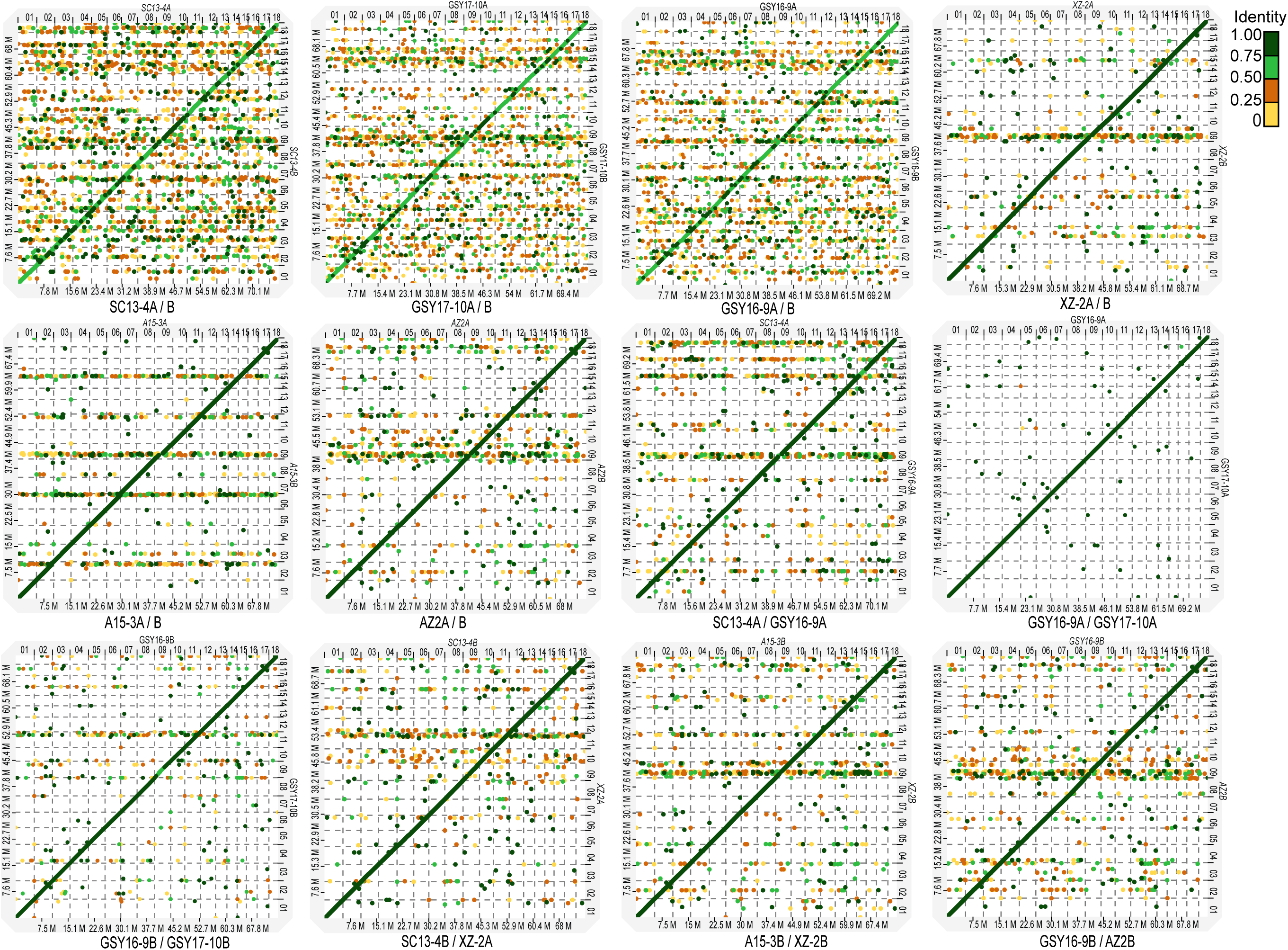
Dot-plot comparisons among haplotype genomes of *Pst* isolates. Dot plots showing pairwise whole-genome alignments between haplotype assemblies from representative *Pst* isolates. Each panel corresponds to a pair of haplotypes, with genomic coordinates displayed on the x and y axes. Points represent aligned regions, and point/line colors indicate sequence identity.

**fig. S6.**
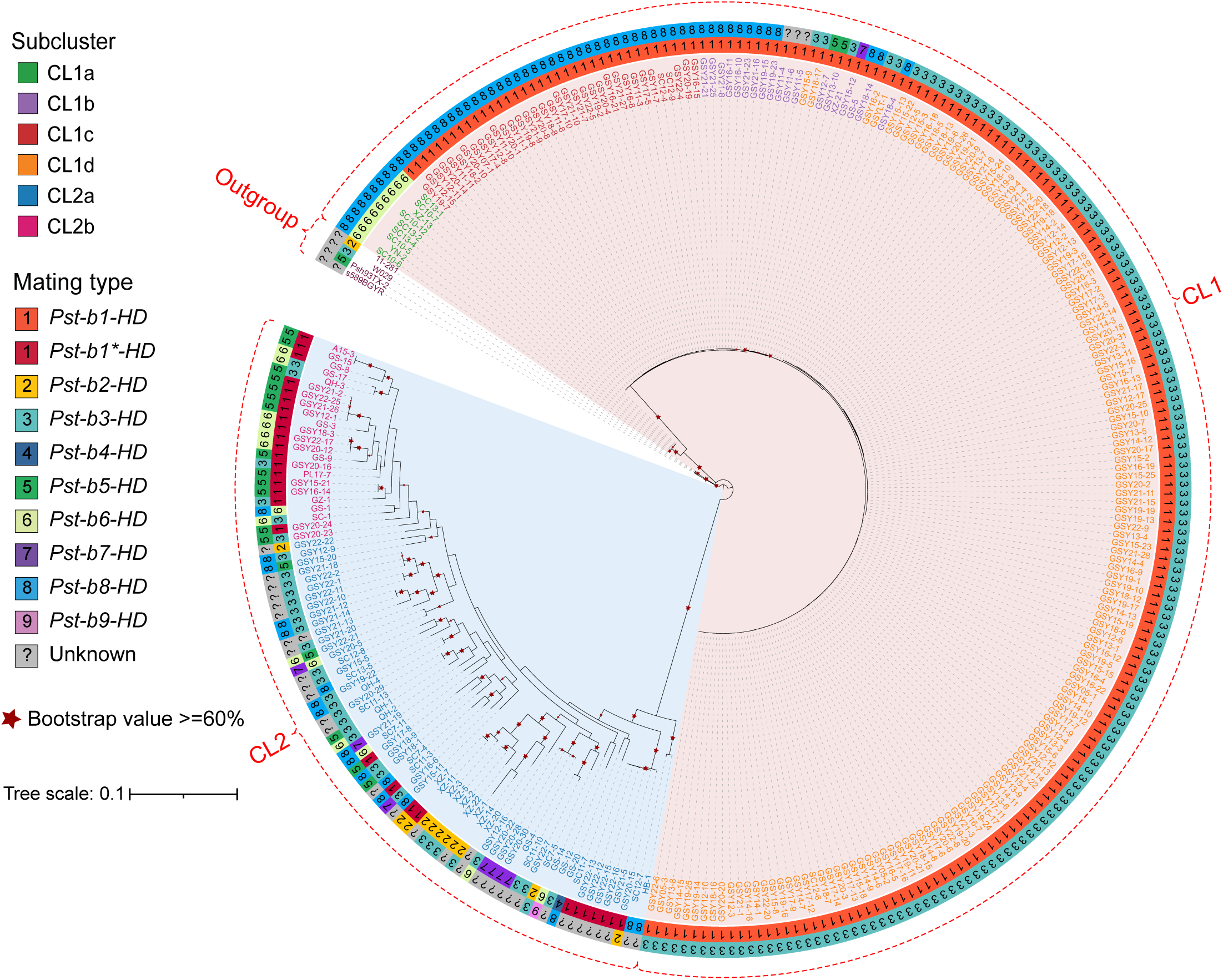
Maximum-likelihood phylogenetic tree of 290 *Pst* isolates constructed using genome-wide SNPs with the GSY16-9A haplotype as the reference. Three *P. striiformis* f. sp. *hordei* isolates and a *P. striiformis* f. sp. *agropyri* isolate are used as outgroups. Subclusters (CL1a to CL1d and CL2a to CL2b) are indicated by colored outer rings. Mating-type alleles (*HD* genes) for each isolate are shown as paired colored bars in the outermost ring; isolates with undetermined mating types are labeled in gray. Red stars denote branches with bootstrap support ≥ 60%.

**fig. S7.**
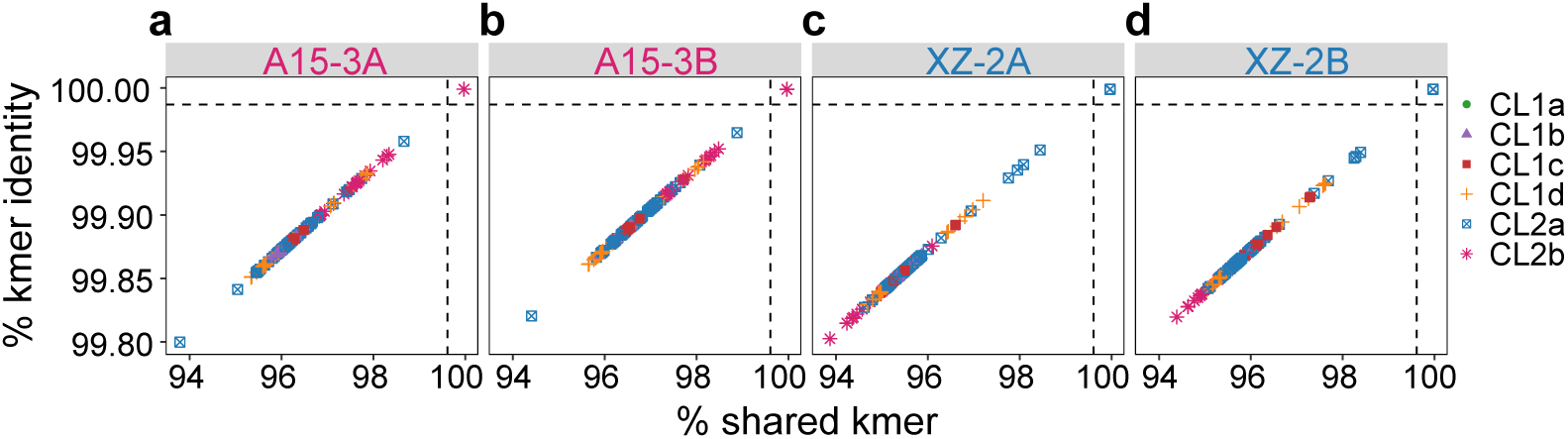
Genome-wide *k*-mer similarity between short-read sequencing data and representative *Pst* reference haplotypes. **a–d**, Scatterplots showing the relationship between the percentage of shared *k*-mers (x axis) and *k*-mer identity (y axis) for 290 isolates relative to four haplotype references: A15-3A **(a)**, A15-3B **(b)**, XZ-2A **(c)** and XZ-2B **(d)**. Each point represents one isolate, colored by its subcluster assignment.

**fig. S8.**
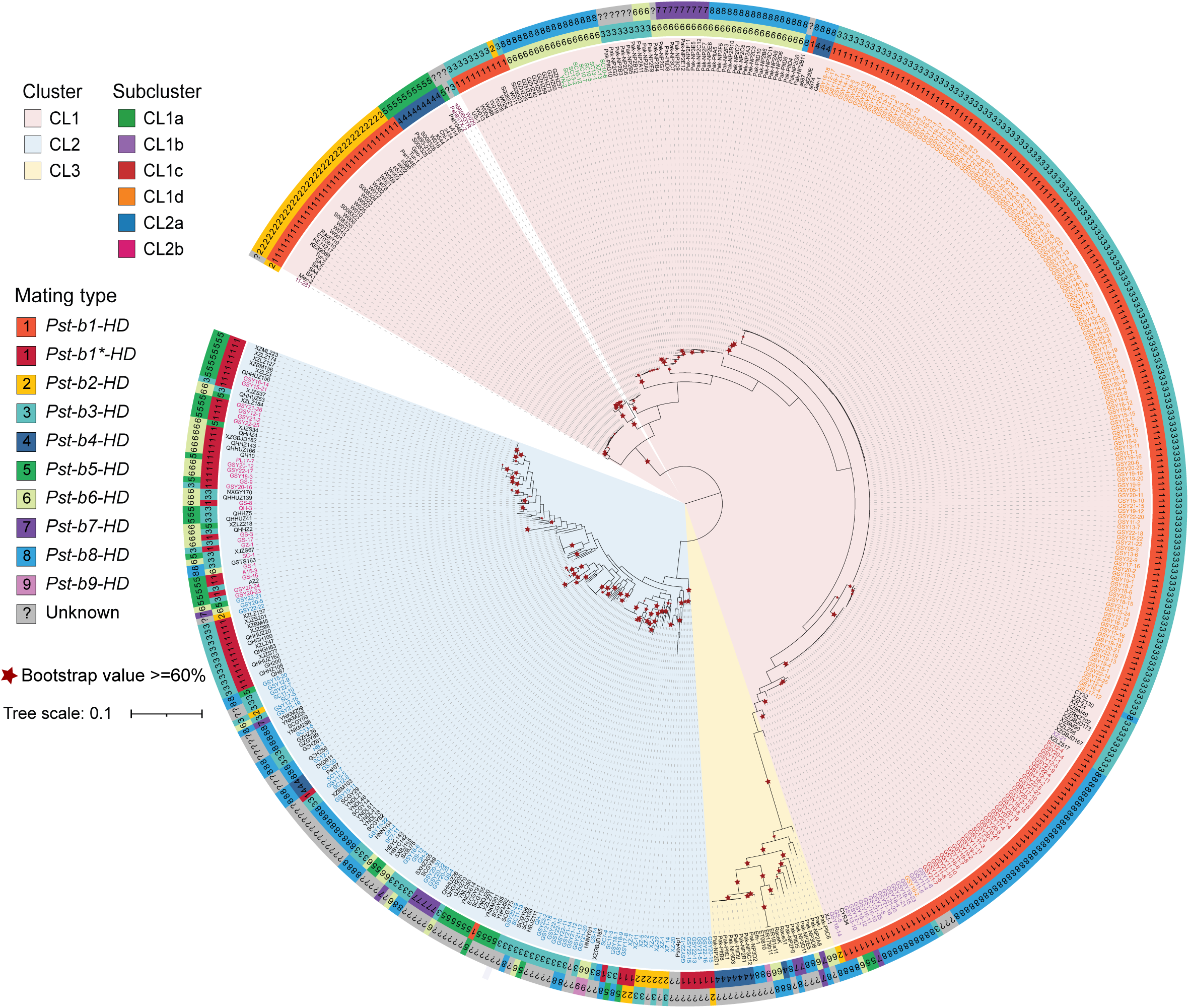
SNP-based phylogeny using the AZ2B haplotype reference recovers the major global *Pst* clusters. Maximum-likelihood phylogenetic tree of global *Pst* isolates constructed from genome-wide SNPs using the AZ2B haplotype as the reference. Isolates are annotated by clusters (CL1–3) and subclusters. Mating-type alleles (*HD* genes) for each isolate are shown as paired colored bars in the outermost ring; isolates with undetermined mating types are labeled in gray. Red stars denote branches with bootstrap support ≥ 60%.

**fig. S9.**
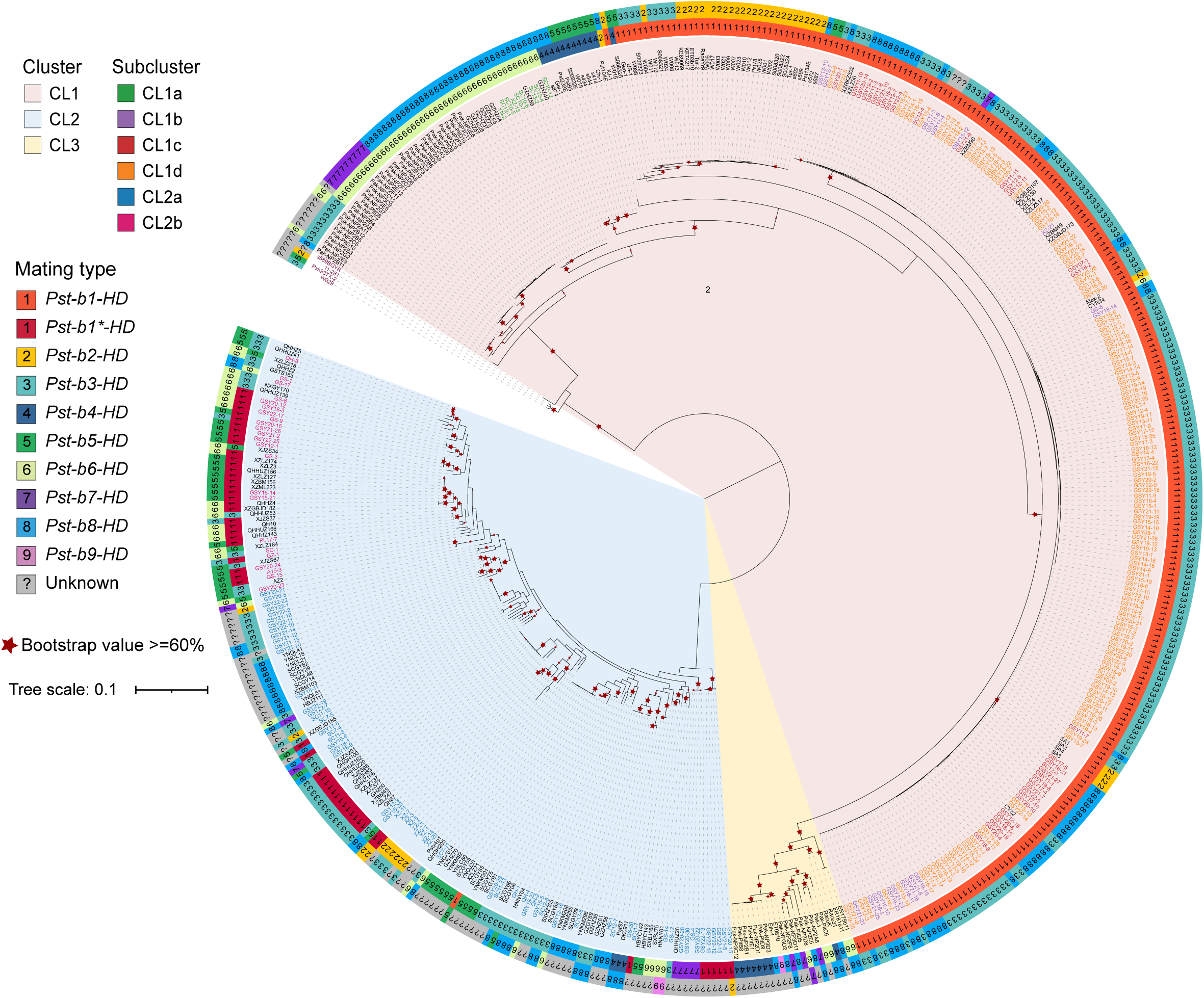
SNP-based phylogeny using the GSY16-9A haplotype reference recovers the major global *Pst* clusters. Maximum-likelihood phylogenetic tree of global *Pst* isolates constructed from genome-wide SNPs using the GSY16-9A haplotype as the reference. Isolates are annotated by clusters (CL1–3) and subclusters. Mating-type alleles (*HD* genes) for each isolate are shown as paired colored bars in the outermost ring; isolates with undetermined mating types are labeled in gray. Red stars denote branches with bootstrap support ≥ 60%.

**fig. S10.**
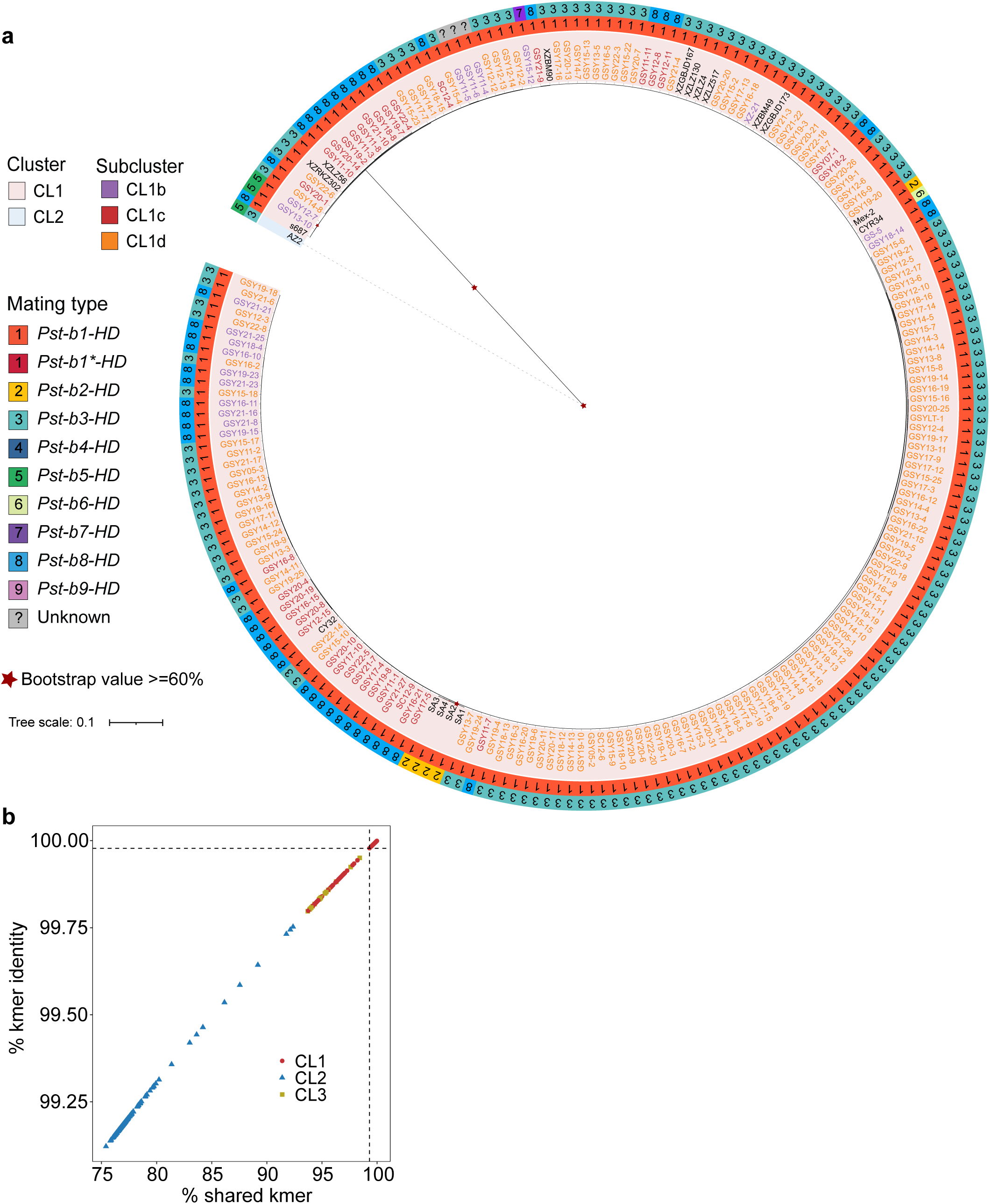
Subclade phylogeny and *k*-mer similarity analysis of global *Pst* isolates. **a**, Subtree extracted from the maximum-likelihood phylogeny constructed from genome-wide SNPs using the GSY16-9A haplotype as the reference. Isolates are annotated by clusters and subclusters. Mating-type alleles (*HD* gene pairs) are shown as paired colored bars in the outermost ring, with gray indicating undetermined mating types. Red stars denote nodes with bootstrap support ≥ 60%. **b**, Scatterplot of percent shared *k*-mers (x axis) versus *k*-mer identity (y axis) comparing isolate short-read data with the GSY16-9A haplotype.

**fig. S11.**
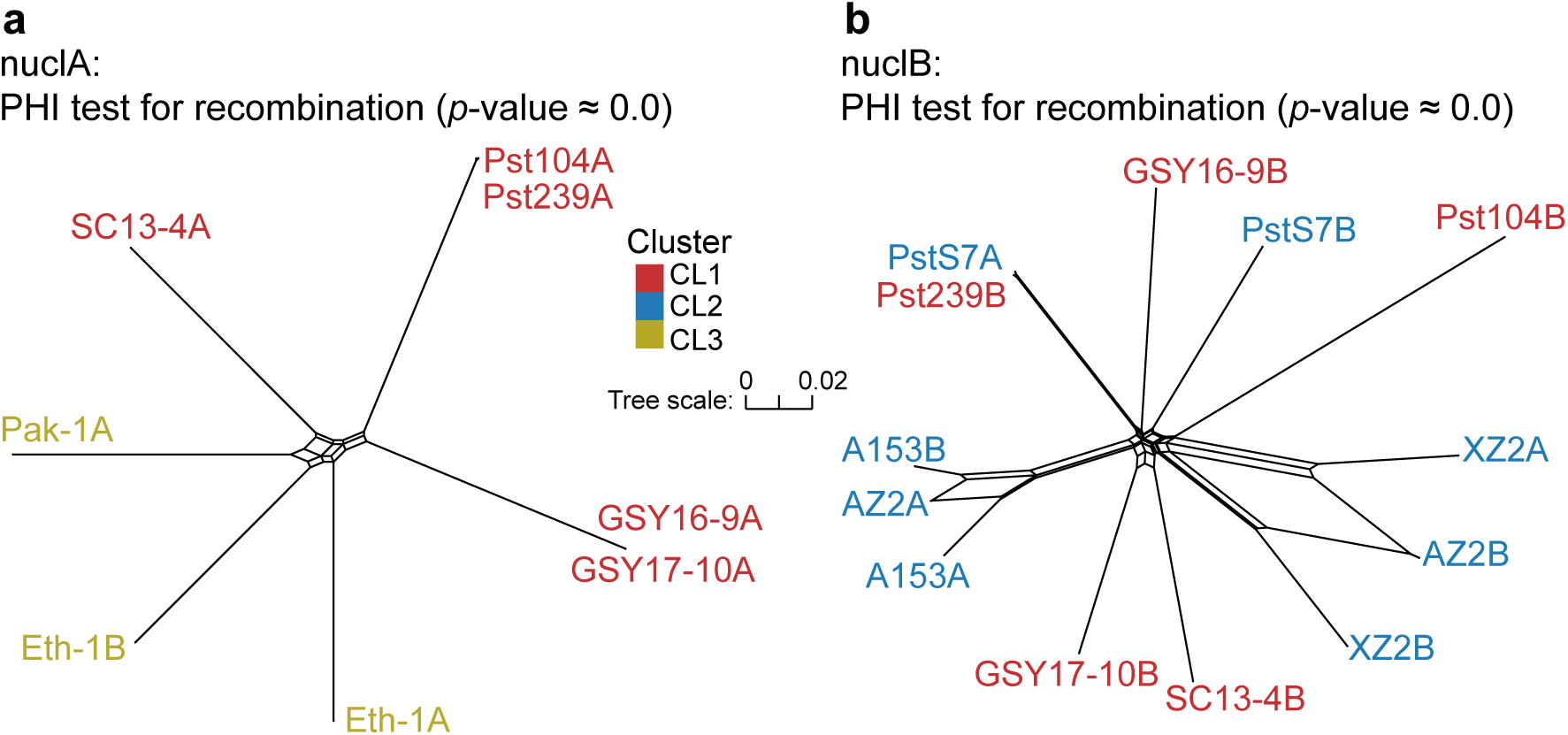
Neighbor-net networks of the two nuclear lineages of *Pst*. **a**, Neighbor-net network of nuclA haplotypes based on genome-wide SNPs. Reticulate structures indicate signals of recombination, supported by the pairwise homoplasy index (PHI) test (*p*-value ≈ 0.0). **b**, Neighbor-net network of nuclB haplotypes. Extensive reticulation similarly reflects intra-lineage recombination, with PHI test *p*-value ≈ 0.0. Colors denote the cluster origin (CL1–3) of each haplotype.

## Supplementary Tables

**Table S1.** Detailed information on *Puccinia striiformis* f. sp. *tritici* isolates used in this study.

| Isolate | Year | Location | Total clean reads | Mapped reads | Mapping rate (%) | Mapping quality | Depth (×) | coverRef 1× (%) | NCBI Accession number |
| --- | --- | --- | --- | --- | --- | --- | --- | --- | --- |
| A15-3 | 2017 | Gansu, China | 80,038,810 | 78,248,851 | 97.76 | 52.97 | 151.51 | 98.21 | SRR26345460 |
| GS-1 | 2016 | Gansu, China | 45,201,538 | 42,392,685 | 93.79 | 52.99 | 81.91 | 98.41 | SRR27320894 |
| GS-12 | 2017 | Gansu, China | 41,606,574 | 35,941,932 | 86.39 | 53.12 | 69.51 | 98.46 | SRR27320857 |
| GS-14 | 2017 | Gansu, China | 44,836,562 | 33,011,889 | 73.63 | 53.05 | 63.80 | 98.96 | SRR27320856 |
| GS-15 | 2017 | Gansu, China | 44,571,283 | 30,663,455 | 68.80 | 53.09 | 59.28 | 98.34 | SRR27320893 |
| GS-17 | 2017 | Gansu, China | 37,704,899 | 20,368,933 | 54.02 | 53.30 | 39.39 | 98.21 | SRR27320892 |
| GS-20 | 2014 | Gansu, China | 52,361,053 | 44,994,851 | 85.93 | 53.29 | 87.06 | 99.62 | SRR27320882 |
| GS-3 | 2014 | Gansu, China | 48,106,559 | 45,697,630 | 94.99 | 53.15 | 88.37 | 98.59 | SRR27320891 |
| GS-4 | 2015 | Gansu, China | 46,867,376 | 45,279,495 | 96.61 | 53.12 | 87.54 | 98.71 | SRR27320890 |
| GS-5 | 2017 | Gansu, China | 46,604,171 | 31,074,770 | 66.68 | 52.28 | 59.42 | 99.06 | SRR27320889 |
| GS-8 | 2015 | Gansu, China | 38,321,631 | 36,658,407 | 95.66 | 53.18 | 70.96 | 98.40 | SRR27320888 |
| GS-9 | 2017 | Gansu, China | 38,345,391 | 36,531,485 | 95.27 | 53.14 | 70.54 | 98.21 | SRR27320887 |
| GSY05-1 | 2005 | Gansu, China | 28,383,670 | 20,021,414 | 70.54 | 52.01 | 38.40 | 98.67 | This study |
| GSY05-2 | 2005 | Gansu, China | 29,329,426 | 26,902,523 | 91.73 | 52.04 | 51.79 | 98.76 | This study |
| GSY05-3 | 2005 | Gansu, China | 30,421,040 | 27,554,095 | 90.58 | 52.11 | 53.13 | 98.74 | This study |
| GSY07-1 | 2007 | Gansu, China | 29,237,438 | 25,363,381 | 86.75 | 51.95 | 48.74 | 98.95 | This study |
| GSY11-1 | 2011 | Gansu, China | 32,995,429 | 16,459,287 | 49.88 | 52.07 | 31.61 | 98.83 | This study |
| GSY11-10 | 2011 | Gansu, China | 32,699,757 | 16,482,389 | 50.41 | 51.82 | 31.58 | 98.84 | This study |
| GSY11-11 | 2011 | Gansu, China | 27,520,066 | 23,511,657 | 85.43 | 51.98 | 45.24 | 98.91 | This study |
| GSY11-2 | 2011 | Gansu, China | 29,878,542 | 16,281,071 | 54.49 | 52.00 | 31.15 | 98.61 | This study |
| GSY11-3 | 2011 | Gansu, China | 31,085,336 | 14,573,456 | 46.88 | 51.88 | 27.91 | 98.83 | This study |
| GSY11-4 | 2011 | Gansu, China | 33,914,750 | 16,606,152 | 48.96 | 52.03 | 31.89 | 98.78 | This study |
| GSY11-5 | 2011 | Gansu, China | 34,393,363 | 10,878,235 | 31.63 | 52.04 | 20.81 | 98.47 | This study |
| GSY11-6 | 2011 | Gansu, China | 30,282,139 | 12,689,582 | 41.90 | 52.04 | 24.33 | 98.58 | This study |
| GSY11-7 | 2011 | Gansu, China | 31,416,013 | 21,817,187 | 69.45 | 52.14 | 42.00 | 98.91 | This study |
| GSY11-8 | 2011 | Gansu, China | 32,417,650 | 9,293,455 | 28.67 | 51.84 | 17.73 | 98.62 | This study |
| GSY11-9 | 2011 | Gansu, China | 35,098,227 | 15,636,713 | 44.55 | 51.86 | 29.94 | 98.62 | This study |
| GSY12-1 | 2012 | Gansu, China | 29,587,617 | 26,682,047 | 90.18 | 53.24 | 51.78 | 97.99 | This study |
| GSY12-10 | 2012 | Gansu, China | 34,408,890 | 15,109,418 | 43.91 | 51.98 | 28.77 | 98.62 | This study |
| GSY12-11 | 2012 | Gansu, China | 27,715,622 | 24,394,013 | 88.02 | 52.19 | 46.97 | 98.94 | This study |
| GSY12-12 | 2012 | Gansu, China | 32,647,614 | 21,159,514 | 64.81 | 52.19 | 40.64 | 98.70 | This study |
| GSY12-13 | 2012 | Gansu, China | 32,401,293 | 18,218,253 | 56.23 | 52.08 | 34.97 | 98.66 | This study |
| GSY12-14 | 2012 | Gansu, China | 26,741,352 | 23,355,620 | 87.34 | 52.08 | 44.93 | 98.70 | This study |
| GSY12-15 | 2012 | Gansu, China | 31,469,185 | 24,063,744 | 76.47 | 52.09 | 46.28 | 98.94 | This study |
| GSY12-16 | 2012 | Gansu, China | 30,285,353 | 23,332,104 | 77.04 | 53.14 | 45.39 | 98.52 | This study |
| GSY12-17 | 2012 | Gansu, China | 27,629,534 | 22,027,978 | 79.73 | 52.13 | 42.36 | 99.20 | This study |
| GSY12-2 | 2012 | Gansu, China | 30,890,411 | 19,152,317 | 62.00 | 51.92 | 36.64 | 98.69 | This study |
| GSY12-3 | 2012 | Gansu, China | 28,523,249 | 18,127,536 | 63.55 | 52.03 | 34.70 | 98.66 | This study |
| GSY12-4 | 2012 | Gansu, China | 32,361,702 | 19,051,926 | 58.87 | 51.86 | 36.50 | 98.67 | This study |
| GSY12-5 | 2012 | Gansu, China | 28,301,882 | 23,618,567 | 83.45 | 51.97 | 45.28 | 98.74 | This study |
| GSY12-6 | 2012 | Gansu, China | 31,258,211 | 27,732,370 | 88.72 | 52.14 | 53.40 | 98.76 | This study |
| GSY12-7 | 2012 | Gansu, China | 32,620,283 | 22,254,425 | 68.22 | 51.92 | 42.73 | 98.99 | This study |
| GSY12-8 | 2012 | Gansu, China | 28,703,473 | 24,423,823 | 85.09 | 52.06 | 46.99 | 98.93 | This study |
| GSY12-9 | 2012 | Gansu, China | 32,124,384 | 29,712,601 | 92.49 | 53.05 | 57.74 | 98.42 | This study |
| GSY13-1 | 2013 | Gansu, China | 28,386,993 | 21,496,297 | 75.73 | 52.19 | 41.32 | 98.71 | This study |
| GSY13-10 | 2013 | Gansu, China | 28,069,642 | 25,254,063 | 89.97 | 52.16 | 48.57 | 99.02 | This study |
| GSY13-11 | 2013 | Gansu, China | 30,912,221 | 23,837,048 | 77.11 | 51.97 | 45.78 | 98.70 | This study |
| GSY13-3 | 2013 | Gansu, China | 28,360,628 | 19,535,915 | 68.88 | 51.93 | 37.54 | 98.65 | This study |
| GSY13-4 | 2013 | Gansu, China | 27,792,769 | 19,834,745 | 71.37 | 52.04 | 38.13 | 98.60 | This study |
| GSY13-5 | 2013 | Gansu, China | 29,469,728 | 28,074,277 | 95.26 | 52.14 | 54.05 | 98.75 | This study |
| GSY13-6 | 2013 | Gansu, China | 33,362,731 | 16,324,393 | 48.93 | 52.08 | 31.32 | 98.62 | This study |
| GSY13-7 | 2013 | Gansu, China | 32,926,645 | 17,720,806 | 53.82 | 52.01 | 34.01 | 98.64 | This study |
| GSY13-8 | 2013 | Gansu, China | 29,082,303 | 15,532,290 | 53.41 | 51.97 | 29.73 | 98.59 | This study |
| GSY13-9 | 2013 | Gansu, China | 28,346,345 | 14,413,353 | 50.85 | 51.96 | 27.57 | 98.59 | This study |
| GSY14-1 | 2014 | Gansu, China | 29,886,524 | 28,631,797 | 95.80 | 52.08 | 55.14 | 98.76 | This study |
| GSY14-10 | 2014 | Gansu, China | 28,712,692 | 16,778,362 | 58.44 | 52.06 | 32.20 | 98.63 | This study |
| GSY14-11 | 2014 | Gansu, China | 31,477,890 | 23,110,238 | 73.42 | 52.12 | 44.45 | 98.70 | This study |
| GSY14-12 | 2014 | Gansu, China | 32,726,783 | 22,794,929 | 69.65 | 52.05 | 43.79 | 98.71 | This study |
| GSY14-13 | 2014 | Gansu, China | 31,522,887 | 23,113,639 | 73.32 | 52.08 | 44.45 | 99.32 | This study |
| GSY14-14 | 2014 | Gansu, China | 33,076,941 | 24,797,002 | 74.97 | 52.12 | 47.70 | 98.74 | This study |
| GSY14-15 | 2014 | Gansu, China | 32,525,706 | 14,120,839 | 43.41 | 52.00 | 27.07 | 98.54 | This study |
| GSY14-16 | 2014 | Gansu, China | 31,229,434 | 24,746,749 | 79.24 | 52.08 | 47.60 | 98.73 | This study |
| GSY14-2 | 2014 | Gansu, China | 33,266,828 | 14,444,075 | 43.42 | 51.82 | 27.57 | 98.60 | This study |
| GSY14-3 | 2014 | Gansu, China | 28,293,727 | 16,774,965 | 59.29 | 51.85 | 32.19 | 98.62 | This study |
| GSY14-4 | 2014 | Gansu, China | 33,641,127 | 8,459,690 | 25.15 | 52.01 | 15.99 | 98.38 | This study |
| GSY14-5 | 2014 | Gansu, China | 30,231,540 | 13,327,365 | 44.08 | 52.04 | 25.52 | 98.55 | This study |
| GSY14-6 | 2014 | Gansu, China | 30,456,827 | 19,063,888 | 62.59 | 52.04 | 36.62 | 98.67 | This study |
| GSY14-7 | 2014 | Gansu, China | 30,987,322 | 17,314,764 | 55.88 | 52.02 | 32.99 | 98.67 | This study |
| GSY14-8 | 2014 | Gansu, China | 28,310,508 | 18,046,003 | 63.74 | 52.04 | 34.59 | 98.66 | This study |
| GSY14-9 | 2014 | Gansu, China | 32,453,011 | 12,718,890 | 39.19 | 51.91 | 24.20 | 98.59 | This study |
| GSY15-1 | 2015 | Gansu, China | 30,098,629 | 26,547,744 | 88.20 | 52.13 | 51.11 | 98.74 | This study |
| GSY15-10 | 2015 | Gansu, China | 30,330,322 | 26,162,163 | 86.26 | 52.14 | 50.35 | 99.39 | This study |
| GSY15-11 | 2015 | Gansu, China | 28,706,634 | 26,188,329 | 91.23 | 53.30 | 50.80 | 98.57 | This study |
| GSY15-12 | 2015 | Gansu, China | 30,939,720 | 28,544,660 | 92.26 | 52.19 | 54.94 | 98.79 | This study |
| GSY15-13 | 2015 | Gansu, China | 28,868,195 | 20,506,772 | 71.04 | 52.09 | 39.39 | 98.68 | This study |
| GSY15-15 | 2015 | Gansu, China | 32,156,332 | 26,018,669 | 80.91 | 52.21 | 50.01 | 98.73 | This study |
| GSY15-16 | 2015 | Gansu, China | 28,447,846 | 17,885,552 | 62.87 | 51.96 | 34.18 | 98.66 | This study |
| GSY15-17 | 2015 | Gansu, China | 33,594,014 | 16,337,128 | 48.63 | 51.87 | 31.23 | 98.60 | This study |
| GSY15-18 | 2015 | Gansu, China | 32,319,523 | 16,213,511 | 50.17 | 52.10 | 31.00 | 98.63 | This study |
| GSY15-19 | 2015 | Gansu, China | 30,198,567 | 15,410,268 | 51.03 | 52.11 | 29.44 | 98.63 | This study |
| GSY15-2 | 2015 | Gansu, China | 30,360,786 | 25,229,940 | 83.10 | 52.15 | 48.56 | 98.81 | This study |
| GSY15-20 | 2015 | Gansu, China | 30,525,184 | 26,645,716 | 87.29 | 53.24 | 51.81 | 98.79 | This study |
| GSY15-21 | 2015 | Gansu, China | 31,307,650 | 26,485,625 | 84.60 | 53.30 | 51.50 | 98.44 | This study |
| GSY15-22 | 2015 | Gansu, China | 31,725,880 | 25,300,517 | 79.75 | 52.14 | 48.69 | 98.74 | This study |
| GSY15-23 | 2015 | Gansu, China | 37,710,995 | 18,530,800 | 49.14 | 52.02 | 35.42 | 98.66 | This study |
| GSY15-24 | 2015 | Gansu, China | 27,827,047 | 22,197,700 | 79.77 | 52.08 | 42.74 | 98.70 | This study |
| GSY15-25 | 2015 | Gansu, China | 33,299,121 | 20,543,711 | 61.69 | 52.01 | 39.19 | 98.71 | This study |
| GSY15-3 | 2015 | Gansu, China | 31,495,312 | 23,377,541 | 74.23 | 52.16 | 44.92 | 98.72 | This study |
| GSY15-4 | 2015 | Gansu, China | 30,265,693 | 15,489,770 | 51.18 | 52.06 | 29.65 | 98.60 | This study |
| GSY15-5 | 2015 | Gansu, China | 29,500,077 | 26,394,884 | 89.47 | 53.09 | 51.29 | 98.87 | This study |
| GSY15-6 | 2015 | Gansu, China | 29,936,887 | 27,691,362 | 92.50 | 52.17 | 53.30 | 98.73 | This study |
| GSY15-7 | 2015 | Gansu, China | 31,475,917 | 10,290,658 | 32.69 | 51.99 | 19.57 | 98.50 | This study |
| GSY15-8 | 2015 | Gansu, China | 31,533,159 | 20,702,590 | 65.65 | 52.12 | 39.76 | 98.70 | This study |
| GSY15-9 | 2015 | Gansu, China | 30,582,929 | 27,933,638 | 91.34 | 52.08 | 53.76 | 98.74 | This study |
| GSY16-10 | 2016 | Gansu, China | 30,313,814 | 27,497,065 | 90.71 | 52.09 | 52.88 | 98.97 | This study |
| GSY16-11 | 2016 | Gansu, China | 29,569,744 | 24,345,604 | 82.33 | 52.21 | 46.85 | 98.95 | This study |
| GSY16-12 | 2016 | Gansu, China | 28,582,031 | 15,305,846 | 53.55 | 52.11 | 29.33 | 98.62 | This study |
| GSY16-13 | 2016 | Gansu, China | 39,560,501 | 13,894,663 | 35.12 | 51.91 | 26.61 | 98.50 | This study |
| GSY16-14 | 2016 | Gansu, China | 39,760,917 | 22,188,372 | 55.80 | 52.95 | 43.07 | 98.50 | This study |
| GSY16-15 | 2016 | Gansu, China | 33,987,945 | 14,732,134 | 43.35 | 52.01 | 28.15 | 98.89 | This study |
| GSY16-18 | 2016 | Gansu, China | 29,661,853 | 17,060,135 | 57.52 | 51.85 | 32.62 | 99.37 | This study |
| GSY16-19 | 2016 | Gansu, China | 32,639,911 | 12,317,080 | 37.74 | 51.99 | 23.47 | 98.58 | This study |
| GSY16-2 | 2016 | Gansu, China | 26,336,572 | 24,720,735 | 93.86 | 52.20 | 47.64 | 99.40 | This study |
| GSY16-20 | 2016 | Gansu, China | 27,676,809 | 21,922,202 | 79.21 | 51.99 | 41.95 | 98.74 | This study |
| GSY16-21 | 2016 | Gansu, China | 27,178,636 | 23,687,900 | 87.16 | 51.97 | 45.52 | 98.94 | This study |
| GSY16-22 | 2016 | Gansu, China | 30,584,965 | 22,724,370 | 74.30 | 52.16 | 43.64 | 98.73 | This study |
| GSY16-3 | 2016 | Gansu, China | 31,044,583 | 25,634,004 | 82.57 | 52.13 | 49.37 | 98.74 | This study |
| GSY16-4 | 2016 | Gansu, China | 30,791,862 | 18,342,948 | 59.57 | 51.81 | 35.16 | 98.65 | This study |
| GSY16-5 | 2016 | Gansu, China | 27,583,550 | 24,199,720 | 87.73 | 52.18 | 46.56 | 99.24 | This study |
| GSY16-6 | 2016 | Gansu, China | 30,389,835 | 22,696,920 | 74.69 | 53.03 | 43.90 | 98.79 | This study |
| GSY16-7 | 2016 | Gansu, China | 30,534,359 | 24,383,115 | 79.85 | 52.08 | 47.01 | 98.68 | This study |
| GSY16-8 | 2016 | Gansu, China | 32,461,745 | 12,270,695 | 37.80 | 52.03 | 23.40 | 98.93 | This study |
| GSY16-9 | 2016 | Gansu, China | 30,645,621 | 29,926,555 | 97.65 | 52.21 | 57.72 | 98.75 | This study |
| GSY17-10 | 2017 | Gansu, China | 36,394,878 | 31,885,668 | 87.61 | 52.03 | 61.38 | 98.98 | This study |
| GSY17-11 | 2017 | Gansu, China | 33,601,736 | 16,488,682 | 49.07 | 51.90 | 31.33 | 98.66 | This study |
| GSY17-12 | 2017 | Gansu, China | 30,733,706 | 20,361,583 | 66.25 | 51.82 | 39.05 | 98.67 | This study |
| GSY17-13 | 2017 | Gansu, China | 28,070,245 | 21,724,912 | 77.39 | 51.98 | 41.71 | 98.72 | This study |
| GSY17-14 | 2017 | Gansu, China | 36,145,577 | 15,784,967 | 43.67 | 51.94 | 30.24 | 98.59 | This study |
| GSY17-15 | 2017 | Gansu, China | 28,135,690 | 24,910,917 | 88.54 | 52.16 | 47.90 | 98.71 | This study |
| GSY17-16 | 2017 | Gansu, China | 30,515,103 | 24,392,514 | 79.94 | 52.11 | 46.93 | 98.73 | This study |
| GSY17-2 | 2017 | Gansu, China | 34,089,106 | 18,943,035 | 55.57 | 52.03 | 36.24 | 98.66 | This study |
| GSY17-3 | 2017 | Gansu, China | 37,210,712 | 12,448,084 | 33.45 | 51.80 | 23.76 | 98.51 | This study |
| GSY17-4 | 2017 | Gansu, China | 33,368,357 | 20,715,090 | 62.08 | 51.97 | 39.83 | 98.91 | This study |
| GSY17-5 | 2017 | Gansu, China | 27,672,752 | 18,794,594 | 67.92 | 52.31 | 36.10 | 98.88 | This study |
| GSY17-6 | 2017 | Gansu, China | 29,361,038 | 23,079,504 | 78.61 | 52.08 | 44.38 | 98.70 | This study |
| GSY17-7 | 2017 | Gansu, China | 28,251,890 | 20,633,914 | 73.04 | 52.17 | 39.66 | 98.67 | This study |
| GSY17-8 | 2017 | Gansu, China | 32,722,498 | 8,157,722 | 24.93 | 52.98 | 15.61 | 98.26 | This study |
| GSY17-9 | 2017 | Gansu, China | 28,495,497 | 25,832,884 | 90.66 | 52.18 | 49.74 | 98.73 | This study |
| GSY18-1 | 2018 | Gansu, China | 27,802,574 | 26,662,558 | 95.90 | 53.20 | 51.79 | 98.79 | This study |
| GSY18-10 | 2018 | Gansu, China | 26,630,118 | 24,450,724 | 91.82 | 52.16 | 47.19 | 98.67 | This study |
| GSY18-12 | 2018 | Gansu, China | 28,717,624 | 26,670,731 | 92.87 | 51.99 | 51.25 | 98.76 | This study |
| GSY18-13 | 2018 | Gansu, China | 31,000,477 | 23,542,797 | 75.94 | 52.07 | 45.31 | 98.73 | This study |
| GSY18-14 | 2018 | Gansu, China | 30,261,930 | 21,949,114 | 72.53 | 52.00 | 42.21 | 99.04 | This study |
| GSY18-15 | 2018 | Gansu, China | 31,988,169 | 23,313,585 | 72.88 | 52.12 | 44.73 | 98.71 | This study |
| GSY18-16 | 2018 | Gansu, China | 32,720,452 | 20,125,968 | 61.51 | 52.09 | 38.67 | 98.69 | This study |
| GSY18-17 | 2018 | Gansu, China | 30,480,461 | 23,986,299 | 78.69 | 52.06 | 46.09 | 98.74 | This study |
| GSY18-2 | 2018 | Gansu, China | 30,697,494 | 29,686,069 | 96.71 | 52.16 | 57.26 | 99.33 | This study |
| GSY18-3 | 2018 | Gansu, China | 35,399,562 | 16,577,768 | 46.83 | 53.06 | 32.10 | 98.07 | This study |
| GSY18-4 | 2018 | Gansu, China | 28,517,125 | 26,613,184 | 93.32 | 52.14 | 51.20 | 98.96 | This study |
| GSY18-5 | 2018 | Gansu, China | 28,526,849 | 20,746,576 | 72.73 | 52.04 | 39.92 | 98.69 | This study |
| GSY18-6 | 2018 | Gansu, China | 32,911,357 | 19,299,234 | 58.64 | 52.07 | 37.13 | 98.67 | This study |
| GSY18-7 | 2018 | Gansu, China | 29,758,365 | 17,392,182 | 58.44 | 52.06 | 33.30 | 98.68 | This study |
| GSY18-8 | 2018 | Gansu, China | 38,139,928 | 13,668,891 | 35.84 | 51.90 | 25.96 | 99.24 | This study |
| GSY18-9 | 2018 | Gansu, China | 30,552,098 | 26,822,156 | 87.79 | 53.23 | 52.24 | 98.67 | This study |
| GSY19-1 | 2019 | Gansu, China | 32,975,907 | 25,272,416 | 76.64 | 52.04 | 48.50 | 98.76 | This study |
| GSY19-10 | 2019 | Gansu, China | 30,304,045 | 27,671,244 | 91.31 | 52.02 | 53.15 | 98.77 | This study |
| GSY19-11 | 2019 | Gansu, China | 26,362,254 | 25,117,741 | 95.28 | 52.16 | 48.46 | 98.72 | This study |
| GSY19-12 | 2019 | Gansu, China | 30,896,801 | 24,146,920 | 78.15 | 52.08 | 46.48 | 98.97 | This study |
| GSY19-13 | 2019 | Gansu, China | 27,966,233 | 24,478,887 | 87.53 | 51.95 | 47.00 | 98.73 | This study |
| GSY19-14 | 2019 | Gansu, China | 36,735,182 | 17,170,797 | 46.74 | 52.20 | 32.85 | 98.67 | This study |
| GSY19-15 | 2019 | Gansu, China | 30,497,893 | 29,568,061 | 96.95 | 52.20 | 57.05 | 98.97 | This study |
| GSY19-16 | 2019 | Gansu, China | 31,007,275 | 27,152,163 | 87.57 | 52.15 | 52.32 | 98.75 | This study |
| GSY19-17 | 2019 | Gansu, China | 32,331,526 | 29,800,731 | 92.17 | 52.14 | 57.46 | 98.76 | This study |
| GSY19-18 | 2019 | Gansu, China | 30,785,233 | 29,137,094 | 94.65 | 52.15 | 56.18 | 98.76 | This study |
| GSY19-19 | 2019 | Gansu, China | 31,151,737 | 29,145,601 | 93.56 | 52.11 | 56.17 | 98.76 | This study |
| GSY19-2 | 2019 | Gansu, China | 31,725,996 | 21,807,378 | 68.74 | 51.84 | 41.89 | 98.89 | This study |
| GSY19-20 | 2019 | Gansu, China | 31,605,459 | 25,253,892 | 79.90 | 52.20 | 48.65 | 98.73 | This study |
| GSY19-21 | 2019 | Gansu, China | 30,416,372 | 26,776,244 | 88.03 | 52.03 | 51.44 | 98.75 | This study |
| GSY19-22 | 2019 | Gansu, China | 27,290,706 | 25,365,429 | 92.95 | 53.13 | 49.25 | 98.56 | This study |
| GSY19-23 | 2019 | Gansu, China | 33,679,699 | 20,635,449 | 61.27 | 52.24 | 39.75 | 98.91 | This study |
| GSY19-24 | 2019 | Gansu, China | 29,960,102 | 28,585,530 | 95.41 | 52.18 | 55.12 | 98.76 | This study |
| GSY19-25 | 2019 | Gansu, China | 31,048,958 | 21,872,873 | 70.45 | 52.02 | 42.15 | 98.71 | This study |
| GSY19-3 | 2019 | Gansu, China | 30,622,155 | 26,441,052 | 86.35 | 52.13 | 50.95 | 98.74 | This study |
| GSY19-4 | 2019 | Gansu, China | 34,970,035 | 30,781,853 | 88.02 | 52.00 | 59.14 | 98.78 | This study |
| GSY19-5 | 2019 | Gansu, China | 27,896,124 | 24,683,887 | 88.49 | 52.15 | 47.39 | 98.74 | This study |
| GSY19-6 | 2019 | Gansu, China | 31,628,434 | 29,019,201 | 91.75 | 52.09 | 55.95 | 98.76 | This study |
| GSY19-7 | 2019 | Gansu, China | 28,350,790 | 17,254,699 | 60.86 | 52.24 | 33.02 | 98.86 | This study |
| GSY19-8 | 2019 | Gansu, China | 30,318,183 | 26,282,153 | 86.69 | 52.06 | 50.66 | 98.94 | This study |
| GSY19-9 | 2019 | Gansu, China | 29,698,886 | 22,803,609 | 76.78 | 52.02 | 43.84 | 98.72 | This study |
| GSY20-1 | 2020 | Gansu, China | 35,857,458 | 18,724,415 | 52.22 | 52.05 | 35.82 | 98.90 | This study |
| GSY20-10 | 2020 | Gansu, China | 32,189,237 | 25,496,829 | 79.21 | 52.17 | 49.11 | 98.95 | This study |
| GSY20-11 | 2020 | Gansu, China | 32,908,324 | 27,879,138 | 84.72 | 52.16 | 53.70 | 98.75 | This study |
| GSY20-12 | 2020 | Gansu, China | 32,918,736 | 25,813,789 | 78.42 | 52.82 | 50.16 | 98.15 | This study |
| GSY20-13 | 2020 | Gansu, China | 29,800,608 | 25,791,363 | 86.55 | 52.08 | 49.67 | 98.74 | This study |
| GSY20-14 | 2020 | Gansu, China | 30,463,498 | 26,863,788 | 88.18 | 52.07 | 51.79 | 98.96 | This study |
| GSY20-15 | 2020 | Gansu, China | 29,338,390 | 28,635,393 | 97.60 | 53.24 | 55.78 | 98.58 | This study |
| GSY20-16 | 2020 | Gansu, China | 31,264,714 | 29,139,972 | 93.20 | 53.25 | 56.75 | 98.63 | This study |
| GSY20-17 | 2020 | Gansu, China | 29,600,438 | 27,855,204 | 94.10 | 52.14 | 53.71 | 98.75 | This study |
| GSY20-18 | 2020 | Gansu, China | 34,241,150 | 8,480,035 | 24.77 | 52.02 | 16.07 | 98.40 | This study |
| GSY20-19 | 2020 | Gansu, China | 33,449,381 | 8,613,166 | 25.75 | 51.87 | 16.22 | 98.64 | This study |
| GSY20-2 | 2020 | Gansu, China | 35,607,333 | 18,355,091 | 51.55 | 52.05 | 35.11 | 98.69 | This study |
| GSY20-20 | 2020 | Gansu, China | 30,389,412 | 22,072,038 | 72.63 | 51.99 | 42.40 | 98.81 | This study |
| GSY20-21 | 2020 | Gansu, China | 33,151,451 | 24,235,477 | 73.11 | 52.23 | 46.43 | 98.73 | This study |
| GSY20-22 | 2020 | Gansu, China | 25,957,821 | 25,325,296 | 97.56 | 53.10 | 49.15 | 98.66 | This study |
| GSY20-23 | 2020 | Gansu, China | 29,545,984 | 23,627,465 | 79.97 | 53.43 | 45.96 | 98.91 | This study |
| GSY20-24 | 2020 | Gansu, China | 29,496,168 | 27,639,691 | 93.71 | 53.15 | 53.81 | 98.57 | This study |
| GSY20-25 | 2020 | Gansu, China | 28,252,974 | 18,221,533 | 64.49 | 52.04 | 34.94 | 98.64 | This study |
| GSY20-26 | 2020 | Gansu, China | 29,638,254 | 28,071,222 | 94.71 | 52.13 | 54.14 | 98.74 | This study |
| GSY20-28 | 2020 | Gansu, China | 30,784,468 | 28,648,598 | 93.06 | 53.12 | 55.79 | 98.67 | This study |
| GSY20-29 | 2020 | Gansu, China | 34,062,868 | 8,769,986 | 25.75 | 53.04 | 16.80 | 98.25 | This study |
| GSY20-3 | 2020 | Gansu, China | 35,789,580 | 17,246,079 | 48.19 | 51.99 | 32.90 | 98.66 | This study |
| GSY20-30 | 2020 | Gansu, China | 30,667,128 | 26,024,247 | 84.86 | 53.12 | 50.54 | 98.65 | This study |
| GSY20-31 | 2020 | Gansu, China | 31,565,131 | 20,096,429 | 63.67 | 52.10 | 38.55 | 98.70 | This study |
| GSY20-4 | 2020 | Gansu, China | 37,835,096 | 12,209,250 | 32.27 | 51.96 | 23.16 | 98.81 | This study |
| GSY20-5 | 2020 | Gansu, China | 31,489,221 | 27,904,410 | 88.62 | 53.21 | 54.36 | 98.76 | This study |
| GSY20-6 | 2020 | Gansu, China | 29,952,632 | 23,889,790 | 79.76 | 52.06 | 45.90 | 98.73 | This study |
| GSY20-7 | 2020 | Gansu, China | 29,591,268 | 26,733,488 | 90.34 | 52.15 | 51.59 | 98.72 | This study |
| GSY20-8 | 2020 | Gansu, China | 33,619,271 | 16,895,222 | 50.25 | 52.07 | 32.56 | 98.79 | This study |
| GSY20-9 | 2020 | Gansu, China | 30,343,106 | 24,118,624 | 79.49 | 52.02 | 46.32 | 98.74 | This study |
| GSY21-1 | 2021 | Gansu, China | 36,529,211 | 22,192,015 | 60.75 | 52.17 | 42.50 | 98.72 | This study |
| GSY21-10 | 2021 | Gansu, China | 29,986,354 | 27,484,117 | 91.66 | 52.07 | 53.00 | 98.96 | This study |
| GSY21-11 | 2021 | Gansu, China | 30,744,692 | 25,977,617 | 84.49 | 52.11 | 50.04 | 98.74 | This study |
| GSY21-12 | 2021 | Gansu, China | 33,176,317 | 21,534,005 | 64.91 | 53.25 | 41.75 | 98.59 | This study |
| GSY21-13 | 2021 | Gansu, China | 29,513,298 | 28,926,909 | 98.01 | 53.29 | 56.34 | 98.29 | This study |
| GSY21-14 | 2021 | Gansu, China | 31,682,863 | 24,593,244 | 77.62 | 53.31 | 47.85 | 98.60 | This study |
| GSY21-15 | 2021 | Gansu, China | 30,914,192 | 22,181,433 | 71.75 | 51.96 | 42.59 | 98.72 | This study |
| GSY21-16 | 2021 | Gansu, China | 31,769,466 | 28,826,223 | 90.74 | 52.08 | 55.36 | 98.96 | This study |
| GSY21-17 | 2021 | Gansu, China | 27,703,857 | 19,699,305 | 71.11 | 52.05 | 37.80 | 98.70 | This study |
| GSY21-18 | 2021 | Gansu, China | 37,004,590 | 26,698,769 | 72.15 | 52.98 | 51.77 | 98.64 | This study |
| GSY21-19 | 2021 | Gansu, China | 31,319,238 | 21,423,575 | 68.40 | 53.15 | 41.63 | 98.84 | This study |
| GSY21-2 | 2021 | Gansu, China | 35,531,077 | 25,094,973 | 70.63 | 53.17 | 48.56 | 98.12 | This study |
| GSY21-20 | 2021 | Gansu, China | 30,582,695 | 24,517,318 | 80.17 | 53.10 | 47.58 | 98.68 | This study |
| GSY21-21 | 2021 | Gansu, China | 29,891,582 | 22,158,551 | 74.13 | 52.15 | 42.68 | 99.03 | This study |
| GSY21-22 | 2021 | Gansu, China | 29,916,133 | 21,160,126 | 70.73 | 52.20 | 40.55 | 98.73 | This study |
| GSY21-23 | 2021 | Gansu, China | 30,674,850 | 28,041,724 | 91.42 | 52.21 | 54.11 | 98.96 | This study |
| GSY21-25 | 2021 | Gansu, China | 30,475,825 | 28,105,784 | 92.22 | 52.29 | 54.23 | 98.97 | This study |
| GSY21-26 | 2021 | Gansu, China | 26,621,067 | 24,832,445 | 93.28 | 53.28 | 48.21 | 97.97 | This study |
| GSY21-27 | 2021 | Gansu, China | 29,277,702 | 18,615,464 | 63.58 | 52.03 | 35.67 | 98.90 | This study |
| GSY21-28 | 2021 | Gansu, China | 29,611,456 | 25,619,686 | 86.52 | 52.13 | 49.31 | 98.76 | This study |
| GSY21-3 | 2021 | Gansu, China | 30,201,789 | 19,472,878 | 64.48 | 52.13 | 37.48 | 98.68 | This study |
| GSY21-4 | 2021 | Gansu, China | 31,463,305 | 23,853,231 | 75.81 | 52.25 | 45.89 | 98.74 | This study |
| GSY21-5 | 2021 | Gansu, China | 30,174,956 | 28,635,462 | 94.90 | 53.32 | 55.80 | 98.49 | This study |
| GSY21-6 | 2021 | Gansu, China | 40,452,908 | 9,886,101 | 24.44 | 51.87 | 18.70 | 98.49 | This study |
| GSY21-7 | 2021 | Gansu, China | 30,993,809 | 16,952,707 | 54.70 | 52.02 | 32.64 | 98.86 | This study |
| GSY21-8 | 2021 | Gansu, China | 34,618,435 | 16,338,770 | 47.20 | 52.14 | 31.25 | 98.88 | This study |
| GSY21-9 | 2021 | Gansu, China | 30,165,320 | 28,552,405 | 94.65 | 51.96 | 54.84 | 98.99 | This study |
| GSY22-1 | 2022 | Gansu, China | 36,024,273 | 27,680,011 | 76.84 | 53.02 | 53.69 | 98.77 | This study |
| GSY22-10 | 2022 | Gansu, China | 32,891,892 | 24,311,411 | 73.91 | 53.15 | 47.20 | 98.53 | This study |
| GSY22-11 | 2022 | Gansu, China | 30,778,994 | 23,119,501 | 75.11 | 53.06 | 44.78 | 98.54 | This study |
| GSY22-12 | 2022 | Gansu, China | 26,591,903 | 23,437,613 | 88.14 | 53.46 | 45.45 | 98.45 | This study |
| GSY22-13 | 2022 | Gansu, China | 31,325,376 | 30,132,769 | 96.19 | 53.30 | 58.63 | 98.50 | This study |
| GSY22-14 | 2022 | Gansu, China | 29,683,060 | 22,969,839 | 77.38 | 52.18 | 44.16 | 98.71 | This study |
| GSY22-15 | 2022 | Gansu, China | 39,377,635 | 18,385,507 | 46.69 | 53.25 | 35.57 | 98.42 | This study |
| GSY22-16 | 2022 | Gansu, China | 35,408,288 | 24,729,295 | 69.84 | 53.56 | 48.03 | 98.45 | This study |
| GSY22-17 | 2022 | Gansu, China | 33,254,720 | 24,332,588 | 73.17 | 53.13 | 47.24 | 98.17 | This study |
| GSY22-18 | 2022 | Gansu, China | 28,941,690 | 25,676,455 | 88.72 | 52.05 | 49.34 | 98.74 | This study |
| GSY22-19 | 2022 | Gansu, China | 42,605,696 | 28,343,615 | 66.53 | 52.40 | 54.46 | 98.76 | This study |
| GSY22-2 | 2022 | Gansu, China | 36,141,303 | 15,152,688 | 41.93 | 53.18 | 29.23 | 98.53 | This study |
| GSY22-20 | 2022 | Gansu, China | 29,695,405 | 27,249,791 | 91.76 | 52.21 | 52.41 | 98.75 | This study |
| GSY22-21 | 2022 | Gansu, China | 29,214,197 | 25,880,073 | 88.59 | 53.02 | 50.26 | 98.82 | This study |
| GSY22-22 | 2022 | Gansu, China | 31,460,144 | 28,284,042 | 89.90 | 53.09 | 54.97 | 98.45 | This study |
| GSY22-25 | 2022 | Gansu, China | 33,337,453 | 20,673,536 | 62.01 | 53.06 | 40.00 | 97.97 | This study |
| GSY22-3 | 2022 | Gansu, China | 29,394,372 | 21,592,031 | 73.46 | 52.21 | 41.39 | 98.72 | This study |
| GSY22-4 | 2022 | Gansu, China | 29,072,757 | 23,078,264 | 79.38 | 51.97 | 44.31 | 98.97 | This study |
| GSY22-5 | 2022 | Gansu, China | 30,179,824 | 22,106,166 | 73.25 | 52.14 | 42.43 | 99.00 | This study |
| GSY22-6 | 2022 | Gansu, China | 30,199,412 | 13,952,284 | 46.20 | 52.26 | 26.66 | 99.29 | This study |
| GSY22-7 | 2022 | Gansu, China | 35,809,860 | 31,088,297 | 86.81 | 53.36 | 60.49 | 98.49 | This study |
| GSY22-8 | 2022 | Gansu, China | 25,059,212 | 22,135,769 | 88.33 | 52.44 | 42.56 | 98.68 | This study |
| GSY22-9 | 2022 | Gansu, China | 29,533,321 | 20,218,674 | 68.46 | 52.20 | 38.83 | 98.68 | This study |
| GSYLT-1 | 2014 | Gansu, China | 38,730,329 | 20,589,906 | 53.16 | 51.88 | 39.51 | 98.69 | This study |
| GZ-1 | 2017 | Guizhou, China | 37,971,351 | 37,057,522 | 97.59 | 53.22 | 71.68 | 98.29 | SRR27320886 |
| HB-1 | 2017 | Hubei, China | 35,217,833 | 34,111,019 | 96.86 | 53.19 | 65.93 | 98.79 | SRR27320885 |
| PL17-7 | 2013 | Gansu, China | 102,295,751 | 98,625,921 | 96.41 | 52.28 | 128.79 | 98.86 | SRR27320861 |
| QH-1 | 2015 | Qinghai, China | 38,246,862 | 35,015,221 | 91.55 | 53.20 | 67.88 | 98.64 | SRR27320880 |
| QH-2 | 2016 | Qinghai, China | 42,511,608 | 39,847,058 | 93.73 | 53.11 | 77.15 | 98.75 | SRR27320879 |
| QH-3 | 2016 | Qinghai, China | 39,987,728 | 38,273,132 | 95.71 | 53.23 | 74.26 | 98.66 | SRR27320878 |
| QH-4 | 2016 | Qinghai, China | 35,276,822 | 30,289,493 | 85.86 | 53.10 | 58.70 | 98.56 | SRR27320877 |
| SC-1 | 2017 | Sichuan, China | 35,989,773 | 33,674,478 | 93.57 | 53.11 | 65.22 | 98.27 | SRR27320876 |
| SC10-12 | 2010 | Sichuan, China | 27,194,314 | 24,159,790 | 88.84 | 51.85 | 46.44 | 98.92 | This study |
| SC10-5 | 2010 | Sichuan, China | 28,940,877 | 22,596,239 | 78.08 | 51.76 | 43.32 | 98.92 | This study |
| SC10-6 | 2010 | Sichuan, China | 32,780,241 | 30,388,855 | 92.70 | 52.13 | 57.45 | 98.97 | This study |
| SC10-7 | 2010 | Sichuan, China | 28,485,355 | 21,100,321 | 74.07 | 51.89 | 40.53 | 98.90 | This study |
| SC11-10 | 2011 | Sichuan, China | 31,621,007 | 28,671,856 | 90.67 | 53.38 | 55.78 | 98.80 | This study |
| SC11-13 | 2011 | Sichuan, China | 33,127,678 | 26,289,010 | 79.36 | 53.26 | 50.24 | 98.50 | This study |
| SC11-3 | 2011 | Sichuan, China | 32,552,690 | 29,651,959 | 91.09 | 53.09 | 56.69 | 98.78 | This study |
| SC11-7 | 2011 | Sichuan, China | 28,130,128 | 26,756,521 | 95.12 | 53.08 | 52.01 | 98.61 | This study |
| SC12-4 | 2012 | Sichuan, China | 29,350,460 | 25,149,119 | 85.69 | 52.13 | 48.38 | 98.97 | This study |
| SC12-6 | 2012 | Sichuan, China | 33,710,120 | 29,163,164 | 86.51 | 52.32 | 55.28 | 98.76 | This study |
| SC12-7 | 2012 | Sichuan, China | 24,907,789 | 22,694,019 | 91.11 | 52.67 | 44.04 | 98.63 | This study |
| SC12-8 | 2012 | Sichuan, China | 30,946,797 | 30,043,931 | 97.08 | 53.24 | 57.76 | 98.85 | This study |
| SC12-9 | 2012 | Sichuan, China | 26,888,577 | 22,925,299 | 85.26 | 52.03 | 44.15 | 98.93 | This study |
| SC13-1 | 2013 | Sichuan, China | 28,923,071 | 26,999,130 | 93.35 | 52.12 | 51.99 | 98.96 | This study |
| SC13-2 | 2013 | Sichuan, China | 27,891,050 | 24,220,452 | 86.84 | 51.85 | 46.56 | 98.93 | This study |
| SC13-4 | 2013 | Sichuan, China | 32,363,189 | 31,285,249 | 96.67 | 52.30 | 59.65 | 98.95 | This study |
| SC13-5 | 2013 | Sichuan, China | 35,264,397 | 24,673,049 | 69.97 | 53.09 | 46.55 | 98.53 | This study |
| SC7-11 | 2007 | Sichuan, China | 27,545,706 | 23,411,248 | 84.99 | 53.09 | 45.50 | 98.58 | This study |
| SC7-4 | 2007 | Sichuan, China | 30,635,070 | 24,860,516 | 81.15 | 53.18 | 48.33 | 98.45 | This study |
| SC7-5 | 2007 | Sichuan, China | 25,198,050 | 22,983,206 | 91.21 | 52.91 | 44.68 | 98.90 | This study |
| XZ-1 | 2016 | Tibet, China | 46,901,745 | 42,842,734 | 91.35 | 53.27 | 82.94 | 98.49 | SRR27320867 |
| XZ-11 | 2016 | Tibet, China | 42,501,446 | 36,700,736 | 86.35 | 53.07 | 71.07 | 99.50 | SRR27320870 |
| XZ-13 | 2016 | Tibet, China | 47,224,854 | 36,845,479 | 78.02 | 52.15 | 70.35 | 98.94 | SRR27320869 |
| XZ-14 | 2016 | Tibet, China | 41,774,121 | 39,547,794 | 94.67 | 53.20 | 76.54 | 98.93 | SRR27320868 |
| XZ-2 | 2016 | Tibet, China | 37,709,133 | 35,068,767 | 93.00 | 53.47 | 68.18 | 100.00 | SRR26345461 |
| XZ-20 | 2015 | Tibet, China | 45,195,775 | 39,230,117 | 86.80 | 52.98 | 75.78 | 99.10 | SRR27320873 |
| XZ-21 | 2017 | Tibet, China | 65,379,780 | 52,022,987 | 79.57 | 52.30 | 99.90 | 99.05 | SRR27320884 |
| XZ-22 | 2016 | Tibet, China | 44,775,087 | 34,203,655 | 76.39 | 53.19 | 66.36 | 99.60 | SRR27320872 |
| XZ-3 | 2016 | Tibet, China | 33,762,159 | 31,732,740 | 93.99 | 53.44 | 61.62 | 99.30 | SRR27320866 |
| XZ-5 | 2016 | Tibet, China | 41,828,980 | 37,066,056 | 88.61 | 53.29 | 72.11 | 100.00 | SRR27320865 |
| XZ-7 | 2017 | Tibet, China | 47,511,893 | 43,473,805 | 91.50 | 53.33 | 84.36 | 99.19 | SRR27320864 |
| YN-2 | 2017 | Yunnan, China | 44,539,289 | 40,180,331 | 90.21 | 52.15 | 77.01 | 98.99 | SRR27320863 |

**Table S2.**
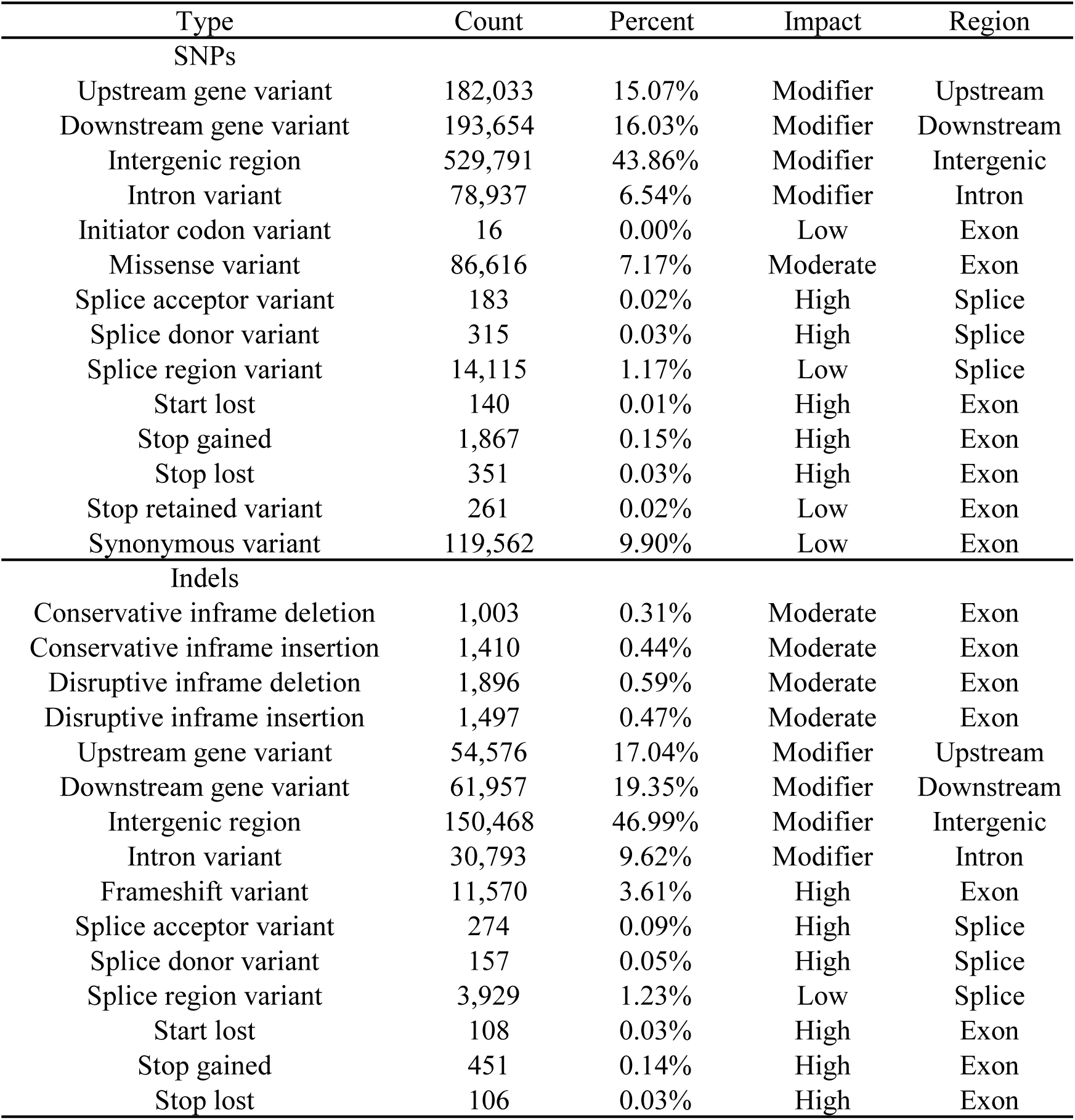
Summary of SNP and indel functional annotations.

| Type | Count | Percent | Impact | Region |
| --- | --- | --- | --- | --- |
| SNPs |  |  |  |  |
| Upstream gene variant | 182,033 | 15.07% | Modifier | Upstream |
| Downstream gene variant | 193,654 | 16.03% | Modifier | Downstream |
| Intergenic region | 529,791 | 43.86% | Modifier | Intergenic |
| Intron variant | 78,937 | 6.54% | Modifier | Intron |
| Initiator codon variant | 16 | 0.00% | Low | Exon |
| Missense variant | 86,616 | 7.17% | Moderate | Exon |
| Splice acceptor variant | 183 | 0.02% | High | Splice |
| Splice donor variant | 315 | 0.03% | High | Splice |
| Splice region variant | 14,115 | 1.17% | Low | Splice |
| Start lost | 140 | 0.01% | High | Exon |
| Stop gained | 1,867 | 0.15% | High | Exon |
| Stop lost | 351 | 0.03% | High | Exon |
| Stop retained variant | 261 | 0.02% | Low | Exon |
| Synonymous variant | 119,562 | 9.90% | Low | Exon |
| Indels |  |  |  |  |
| Conservative inframe deletion | 1,003 | 0.31% | Moderate | Exon |
| Conservative inframe insertion | 1,410 | 0.44% | Moderate | Exon |
| Disruptive inframe deletion | 1,896 | 0.59% | Moderate | Exon |
| Disruptive inframe insertion | 1,497 | 0.47% | Moderate | Exon |
| Upstream gene variant | 54,576 | 17.04% | Modifier | Upstream |
| Downstream gene variant | 61,957 | 19.35% | Modifier | Downstream |
| Intergenic region | 150,468 | 46.99% | Modifier | Intergenic |
| Intron variant | 30,793 | 9.62% | Modifier | Intron |
| Frameshift variant | 11,570 | 3.61% | High | Exon |
| Splice acceptor variant | 274 | 0.09% | High | Splice |
| Splice donor variant | 157 | 0.05% | High | Splice |
| Splice region variant | 3,929 | 1.23% | Low | Splice |
| Start lost | 108 | 0.03% | High | Exon |
| Stop gained | 451 | 0.14% | High | Exon |
| Stop lost | 106 | 0.03% | High | Exon |

**Table S3.**
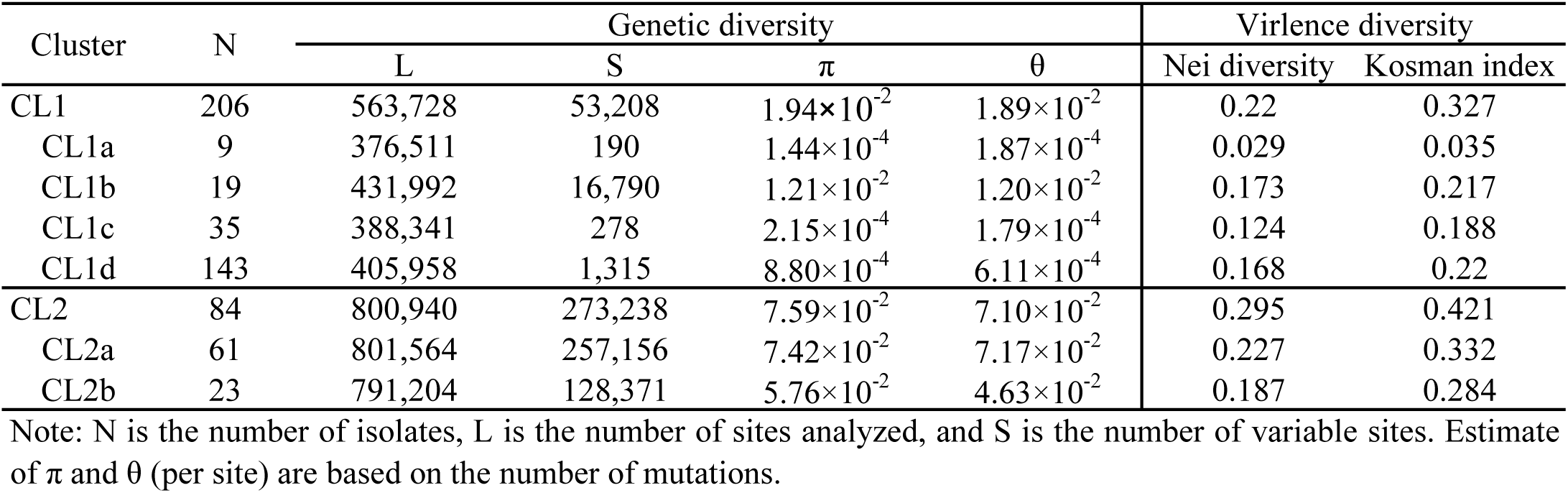
Summary of polymorphisms across genetic clusters of *Puccinia striiformis* f. sp. *tritici*.

**Table S4.**
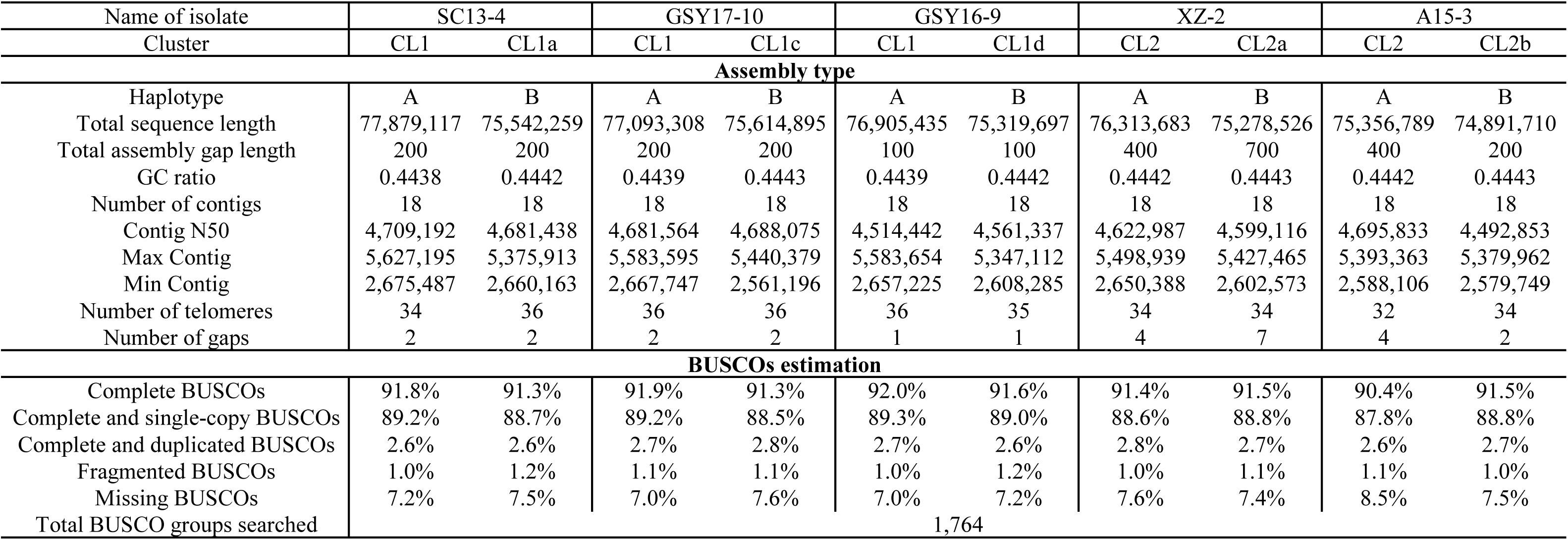
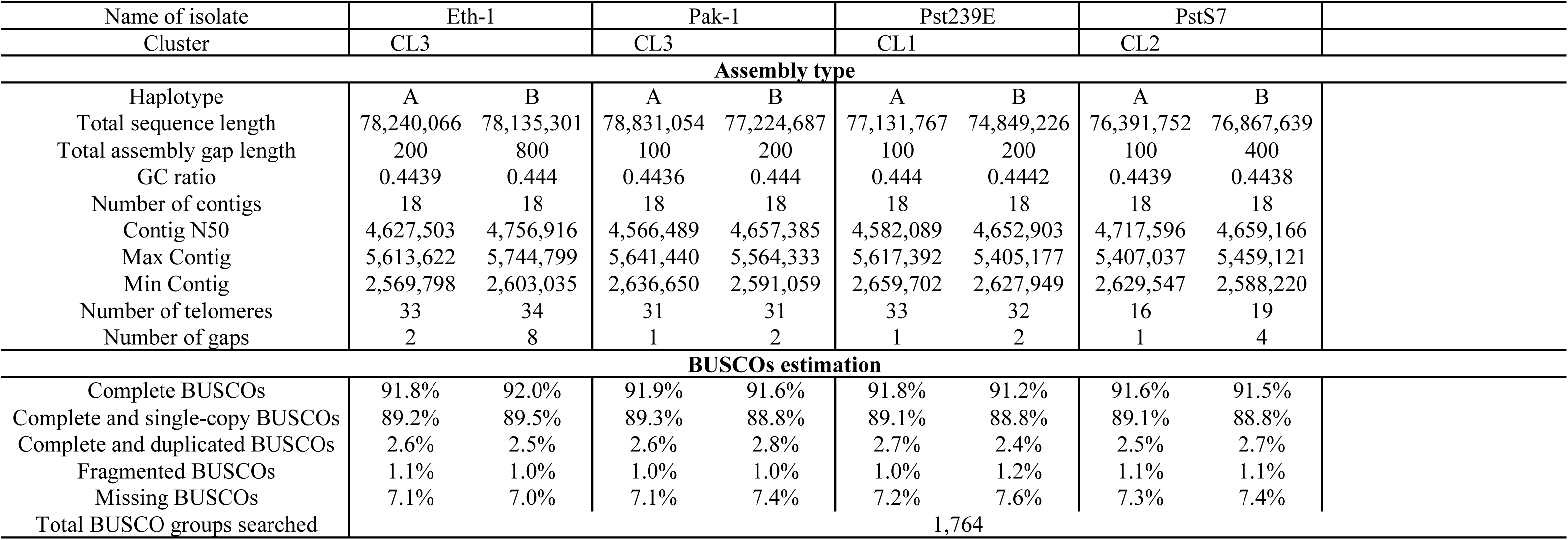
Summary of genome-assembly statistics for *Puccinia striiformis* f. sp. *tritici* isolates.

**Table S5.** Intra- and inter-isolate whole-haplotype alignments for *Puccinia striiformis* f. sp. t*ritici* isolates GSY16-9 and GSY17-10.

|  | GSY16-9A vs GSY16-9B | GSY17-10A vs GSY17-10B | GSY16-9A vs GSY17-10A | GSY16-9B vs GSY17-10B |
| --- | --- | --- | --- | --- |
| Aligned bases | 92.41% | 92.29% | 99.71% | 98.50% |
| Average identity of 1-to-1 alignments | 98.54% | 98.55% | 99.99% | 99.63% |
| Average identity of M-to-M alignments | 98.11% | 98.11% | 99.98% | 99.46% |
| Translocations | 1,292 | 1,312 | 0 | 208 |
| Inversions | 95 | 96 | 0 | 33 |
| Insertions | 12,350 | 12,660 | 250 | 3,640 |
| Total SNPs | 602,125 | 602,558 | 268 | 159,628 |
| Total Indels | 288,479 | 285,869 | 1,985 | 89,357 |

**Table S6.** Detailed information of additional global *Puccinia striiformis* f. sp. *tritici* isolates used in this study.

| Isolate | Year | Location | NCBI Accession number |
| --- | --- | --- | --- |
| GSTS163 | 2015 | Gansu, China | SRR18091585 |
| GZGY89 | 2015 | Guizhou, China | SRR18091528 |
| GZHZ28 | 2015 | Guizhou, China | SRR18091584 |
| GZHZ36 | 2015 | Guizhou, China | SRR18091490 |
| GZHZ37 | 2015 | Guizhou, China | SRR18091492 |
| GZHZ40 | 2015 | Guizhou, China | SRR18091562 |
| GZHZ56 | 2015 | Guizhou, China | SRR18091491 |
| GZHZ57 | 2015 | Guizhou, China | SRR18091483 |
| GZHZ59 | 2015 | Guizhou, China | SRR18091487 |
| GZHZ61 | 2015 | Guizhou, China | SRR18091486 |
| GZHZ64 | 2015 | Guizhou, China | SRR18091519 |
| GZHZ65 | 2015 | Guizhou, China | SRR18091497 |
| GZHZ70 | 2015 | Guizhou, China | SRR18091489 |
| GZHZ73 | 2015 | Guizhou, China | SRR18091508 |
| HBJZ111 | 2015 | Hubei, China | SRR18091481 |
| HBYC142 | 2015 | Hubei, China | SRR18091476 |
| HBYC143 | 2015 | Hubei, China | SRR18091478 |
| HNNY01 | 2015 | Henan, China | SRR18091473 |
| HNNY04 | 2015 | Henan, China | SRR18091470 |
| NXGY170 | 2015 | Ningxia, China | SRR18091461 |
| QH10 | 2015 | Qinghai, China | SRR18091535 |
| QH87 | 2015 | Qinghai, China | SRR18091551 |
| GH200 | 2015 | Qinghai, China | SRR18091549 |
| QHGH100 | 2015 | Qinghai, China | SRR18091552 |
| QHGH205 | 2015 | Qinghai, China | SRR18091554 |
| QHGH83 | 2015 | Qinghai, China | SRR18091555 |
| QHHUZ139 | 2015 | Qinghai, China | SRR18091542 |
| QHHUZ156 | 2015 | Qinghai, China | SRR18091544 |
| QHHUZ162 | 2015 | Qinghai, China | SRR18091541 |
| QHHUZ166 | 2015 | Qinghai, China | SRR18091540 |
| QHHUZ20 | 2015 | Qinghai, China | SRR18091538 |
| QHHUZ26 | 2015 | Qinghai, China | SRR18091537 |
| QHHUZ41 | 2015 | Qinghai, China | SRR18091546 |
| QHHUZ53 | 2015 | Qinghai, China | SRR18091548 |
| QHHZ108 | 2015 | Qinghai, China | SRR18091529 |
| QHHZ143 | 2015 | Qinghai, China | SRR18091532 |
| QHHZ2 | 2015 | Qinghai, China | SRR18091531 |
| QHHZ4 | 2015 | Qinghai, China | SRR18091533 |
| QHHZ5 | 2015 | Qinghai, China | SRR18091534 |
| SCGY06 | 2015 | Sichuan, China | SRR18091458 |
| SCGY09 | 2015 | Sichuan, China | SRR18091526 |
| SCGY14 | 2015 | Sichuan, China | SRR18091457 |
| SCGY29 | 2015 | Sichuan, China | SRR18091525 |
| SCGY55 | 2015 | Sichuan, China | SRR18091450 |
| SCGY65 | 2015 | Sichuan, China | SRR18091451 |
| SCGY66 | 2015 | Sichuan, China | SRR18091448 |
| SCGY75 | 2015 | Sichuan, China | SRR18091444 |
| SCGY82 | 2015 | Sichuan, China | SRR18091447 |
| SCGY89 | 2015 | Sichuan, China | SRR18091453 |
| SCGY91 | 2015 | Sichuan, China | SRR18091446 |
| SXBJ165 | 2015 | Shaanxi, China | SRR18091441 |
| SXBJ75 | 2015 | Shaanxi, China | SRR18091439 |
| SXHZ305 | 2015 | Shaanxi, China | SRR18091436 |
| XJZS201 | 2015 | Xinjiang, China | SRR18091434 |
| XJZS34 | 2015 | Xinjiang, China | SRR18091435 |
| XJZS37 | 2015 | Xinjiang, China | SRR18091582 |
| XJZS67 | 2015 | Xinjiang, China | SRR18091433 |
| XJZS77 | 2015 | Xinjiang, China | SRR18091581 |
| XJZS98 | 2015 | Xinjiang, China | SRR18091583 |
| XZBM103 | 2015 | Xizang, China | SRR18091577 |
| XZBM156 | 2015 | Xizang, China | SRR18091561 |
| XZBM45 | 2015 | Xizang, China | SRR18091564 |
| XZBM49 | 2015 | Xizang, China | SRR18091566 |
| XZBM90 | 2015 | Xizang, China | SRR18091578 |
| XZGBJD167 | 2015 | Xizang, China | SRR18091559 |
| XZGBJD173 | 2015 | Xizang, China | SRR18091560 |
| XZGBJD182 | 2015 | Xizang, China | SRR18091558 |
| XZGBJD185 | 2015 | Xizang, China | SRR18091557 |
| XZLZ127 | 2015 | Xizang, China | SRR18091575 |
| XZLZ130 | 2015 | Xizang, China | SRR18091521 |
| XZLZ137 | 2015 | Xizang, China | SRR18091556 |
| XZLZ174 | 2015 | Xizang, China | SRR18091520 |
| XZLZ184 | 2015 | Xizang, China | SRR18091518 |
| XZLZ218 | 2015 | Xizang, China | SRR18091514 |
| XZLZ3 | 2015 | Xizang, China | SRR18091574 |
| XZLZ47 | 2015 | Xizang, China | SRR18091523 |
| XZLZ4 | 2015 | Xizang, China | SRR18091571 |
| XZLZ517 | 2015 | Xizang, China | SRR18091579 |
| XZLZ56 | 2015 | Xizang, China | SRR18091580 |
| XZLZ71 | 2015 | Xizang, China | SRR18091522 |
| XZML223 | 2015 | Xizang, China | SRR18091517 |
| XZRKZ302 | 2015 | Xizang, China | SRR18091513 |
| YNCX614 | 2015 | Yunnan, China | SRR18091512 |
| YNDL18 | 2015 | Yunnan, China | SRR18091510 |
| YNDL21 | 2015 | Yunnan, China | SRR18091507 |
| YNDL41 | 2015 | Yunnan, China | SRR18091505 |
| YNDL46 | 2015 | Yunnan, China | SRR18091504 |
| YNDL51 | 2015 | Yunnan, China | SRR18091509 |
| YNKM208 | 2015 | Yunnan, China | SRR18091496 |
| YNKM298 | 2015 | Yunnan, China | SRR18091495 |
| YNKM299 | 2015 | Yunnan, China | SRR18091499 |
| YNKM301 | 2015 | Yunnan, China | SRR18091498 |
| YNKM92 | 2015 | Yunnan, China | SRR18091501 |
| YNLC50 | 2015 | Yunnan, China | SRR18091494 |
| YNQJ20 | 2015 | Yunnan, China | SRR18091493 |
| Ger-1 | 2013 | Germany | SRR27320862 |
| Chi-1 | 2014 | Chile | SRR27320895 |
| Geo-1 | 2013 | Georgia | SRR27320858 |
| US-1 | 2013 | USA | SRR27320874 |
| Tur-2 | 2013 | Turkey | SRR27320875 |
| Tur-1 | 2013 | Turkey | SRR27320896 |
| Mex-2 | 2014 | Mexico | SRR27320883 |
| XJ-1 | 2013 | Xinjiang, China | SRR27320871 |
| Pak-1 | 2015 | Pakistan | SRR27320881 |
| Eth-1 | 2015 | Ethiopia | SRR27320860 |
| AZ2 | 2022 | China | SRR26344777 |
| CY32 | NA | China | SRR172667 |
| CYR34 | NA | China | SRR18246449 |
| DK0911 | 2011 | Denmark | SRR10406086 |
| ER179b11 | 2011 | Eritrea | ERR3856811 |
| ER181a11 | 2011 | Eritrea | ERR3856812 |
| ET03b10 | 2010 | Ethiopia | ERR3856810 |
| ET0810 | 2010 | Ethiopia | ERR3856809 |
| KE74217 | 1974 | Kenya | ERR3856807 |
| KE89069 | 1989 | Kenya | ERR3856808 |
| Pst104E | 1982 | Australia | SRR6031720 |
| Pst134E | 2017 | Australia | SRR15357740 |
| Pst78 | 2000 | USA | SRR987408 |
| Pst93-210 | 1993 | USA | SRR8208848 |
| Pstv4p1 | NA | USA | SRR10829480 |
| Race31 | 1936 | India | SRR8601347 |
| RaceK | 1982 | India | SRR8601365 |
| RaceYr9 | 1996 | India | SRR8601595 |
| S008320 | NA | Australia | SRR15496042 |
| S008321 | NA | Australia | SRR15496041 |
| S008322 | NA | Australia | SRR15496040 |
| S008323 | NA | Australia | SRR15496039 |
| S008324 | NA | Australia | SRR15496038 |
| S008325 | NA | Australia | SRR15496037 |
| S008326 | NA | Australia | SRR15496036 |
| s414 | NA | Australia | SRR13781820 |
| s434 | NA | Australia | SRR13781819 |
| s544 | NA | Australia | SRR13799081 |
| s572 | NA | Australia | SRR13799080 |
| s599 | NA | Australia | SRR13799079 |
| s602 | NA | Australia | SRR13799078 |
| s674 | NA | Australia | SRR13823727 |
| s687 | NA | Australia | SRR13823726 |
| SA1 | 1996 | South Africa | ERR3856826 |
| SA2 | 1998 | South Africa | ERR3856827 |
| SA3 | 2001 | South Africa | ERR3856828 |
| SA4 | 2005 | South Africa | ERR3856829 |
| W001 | 2011 | Canada | SRR24010203 |
| W002 | 2011 | Canada | SRR24010202 |
| W003 | 2011 | Canada | SRR24010212 |
| W004 | 2011 | Canada | SRR24010252 |
| W006 | 2011 | Canada | SRR24010211 |
| W007 | 2011 | Canada | SRR24010210 |
| W008 | 2011 | Canada | SRR24010207 |
| W009 | 2011 | Canada | SRR24010204 |
| W010 | 2011 | Canada | SRR24010209 |
| W011 | 2011 | Canada | SRR24010208 |
| W012 | 2011 | Canada | SRR24010206 |
| W013 | 2013 | Canada | SRR24010237 |
| W015 | 2010 | Canada | SRR24010205 |
| W017 | 2010 | Canada | SRR24010201 |
| W018 | 2011 | Canada | SRR24010227 |
| W019 | 2013 | Canada | SRR24010198 |
| W021 | 2005 | Canada | SRR24010200 |
| W023 | 2005 | Canada | SRR24010197 |
| W024 | 2005 | Canada | SRR24010196 |
| W025 | 2005 | Canada | SRR24010199 |
| W088 | 2013 | Canada | SRR24010251 |
| Pst239E | NA | Australia | SRR33364114 |
| PstS7 | NA | NA | SRR33364111 |
| Pak-PltD8 | NA | Pakistan | SRR31145788 |
| Pak-PltD6 | NA | Pakistan | SRR31145789 |
| Pak-PltD5 | NA | Pakistan | SRR31145790 |
| Pak-PltD4 | NA | Pakistan | SRR31145791 |
| Pak-NP3E5 | NA | Pakistan | SRR31145792 |
| Pak-NP3D11 | NA | Pakistan | SRR31145793 |
| Pak-PltA5 | NA | Pakistan | SRR31145794 |
| Pak-NP3D8 | NA | Pakistan | SRR31145795 |
| Pak-NP3D3 | NA | Pakistan | SRR31145796 |
| Pak-NP3D2 | NA | Pakistan | SRR31145797 |
| Pak-NP3C12 | NA | Pakistan | SRR31145798 |
| Pak-NP3C11 | NA | Pakistan | SRR31145799 |
| Pak-NP3C9 | NA | Pakistan | SRR31145800 |
| Pak-NP3C7 | NA | Pakistan | SRR31145801 |
| Pak-NP2G7 | NA | Pakistan | SRR31145802 |
| Pak-NP2G6 | NA | Pakistan | SRR31145803 |
| Pak-NP2G2 | NA | Pakistan | SRR31145804 |
| Pak-PltG10 | NA | Pakistan | SRR31145805 |
| Pak-NP2F11 | NA | Pakistan | SRR31145806 |
| Pak-NP2F8 | NA | Pakistan | SRR31145807 |
| Pak-NP2F7 | NA | Pakistan | SRR31145808 |
| Pak-NP2F5 | NA | Pakistan | SRR31145809 |
| Pak-NP2F3 | NA | Pakistan | SRR31145810 |
| Pak-NP2E12 | NA | Pakistan | SRR31145811 |
| Pak-NP2E9 | NA | Pakistan | SRR31145812 |
| Pak-NP2E6 | NA | Pakistan | SRR31145813 |
| Pak-NP2E5 | NA | Pakistan | SRR31145814 |
| Pak-NP2D6 | NA | Pakistan | SRR31145815 |
| Pak-PltD10 | NA | Pakistan | SRR31145816 |
| Pak-NP2D1 | NA | Pakistan | SRR31145817 |
| Pak-NP2C12 | NA | Pakistan | SRR31145818 |
| Pak-NP2C7 | NA | Pakistan | SRR31145819 |
| Pak-NP2C6 | NA | Pakistan | SRR31145820 |
| Pak-NP2C5 | NA | Pakistan | SRR31145821 |
| Pak-NP2C3 | NA | Pakistan | SRR31145822 |
| Pak-NP2B12 | NA | Pakistan | SRR31145823 |
| Pak-NP2B11 | NA | Pakistan | SRR31145824 |
| Pak-NP2B10 | NA | Pakistan | SRR31145825 |
| Pak-NP2B6 | NA | Pakistan | SRR31145826 |
| Pak-PltD9 | NA | Pakistan | SRR31145827 |
| Pak-NP2B5 | NA | Pakistan | SRR31145828 |
| Pak-NP2B4 | NA | Pakistan | SRR31145829 |
| Pak-NP2B3 | NA | Pakistan | SRR31145830 |
| Pak-NP2B1 | NA | Pakistan | SRR31145831 |
| Pak-NP2A11 | NA | Pakistan | SRR31145832 |
| Pak-NP2A9 | NA | Pakistan | SRR31145833 |
| Pak-NP2A8 | NA | Pakistan | SRR31145834 |
| Pak-NP2A6 | NA | Pakistan | SRR31145835 |
| Pak-NP2A3 | NA | Pakistan | SRR31145836 |
| Pak-PltE1 | NA | Pakistan | SRR31145837 |
| Pak-PltC6 | NA | Pakistan | SRR31145838 |
| Pak-PltB8 | NA | Pakistan | SRR31145839 |

**Table S7.**
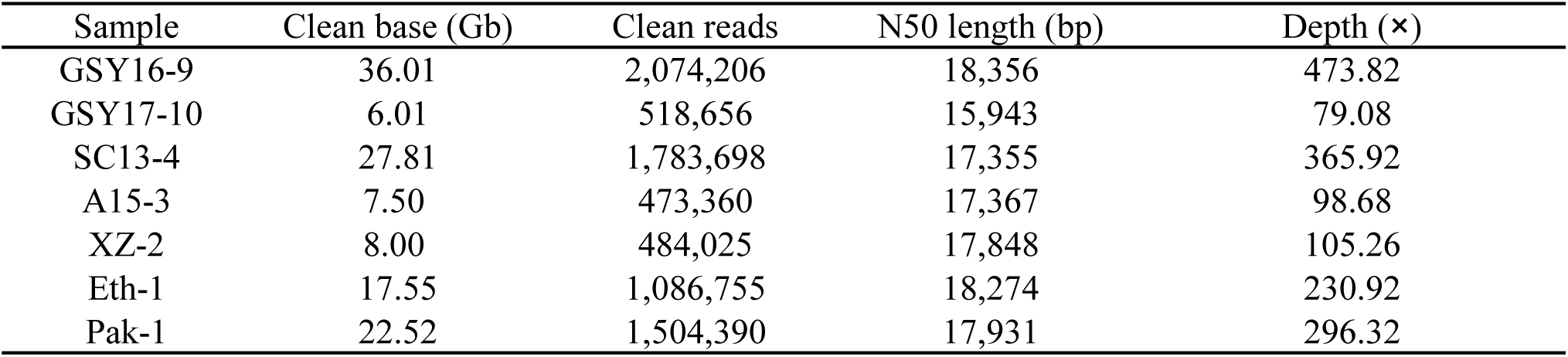
Summary of PacBio HiFi long-read sequencing output.

| Sample | Clean base (Gb) | Clean reads | N50 length (bp) | Depth (×) |
| --- | --- | --- | --- | --- |
| GSY16-9 | 36.01 | 2,074,206 | 18,356 | 473.82 |
| GSY17-10 | 6.01 | 518,656 | 15,943 | 79.08 |
| SC13-4 | 27.81 | 1,783,698 | 17,355 | 365.92 |
| A15-3 | 7.50 | 473,360 | 17,367 | 98.68 |
| XZ-2 | 8.00 | 484,025 | 17,848 | 105.26 |
| Eth-1 | 17.55 | 1,086,755 | 18,274 | 230.92 |
| Pak-1 | 22.52 | 1,504,390 | 17,931 | 296.32 |

**Table S8.** Summary of Hi-C sequencing data.

| Sample | Clean base (Gb) | Clean reads | N50 length (bp) | Depth (×) |
| --- | --- | --- | --- | --- |
| GSY16-9 | 7.82 | 52,161,394 | 2 × 150 | 102.89 |
| GSY17-10 | 9.13 | 60,834,438 | 2 × 150 | 120.13 |
| Eth-1 | 7.38 | 49,180,776 | 2 × 150 | 97.11 |
| Pak-1 | 7.78 | 51,828,221 | 2 × 150 | 102.37 |

